# An *in vivo* CRISPR screen unveils top target genes to improve CAR-T cell efficacy in a solid tumor model

**DOI:** 10.1101/2025.03.28.645780

**Authors:** Mattia Fumagalli, Dongjie An, Luca Simula, Camille Combe, Lisa Aziez, Yannick Simoni, Marie-Clotilde Alves-Guerra, Andrea Valentini, Maude Marchais, Anaïs Vermare, Josquin Moraly, Nadege Bercovici, Emmanuel Donnadieu, Frederic Pendino

## Abstract

CAR-T cell therapies are revolutionizing the treatment of refractory and relapsed haematological malignancies, but many patients do not exhibit long-term responses, and these therapies are less effective against solid tumors. Poor persistence of CAR-T cells in patients is associated with therapeutic failure, highlighting the need to identify strategies promoting *in vivo* expansion. Here, we developed an *in vivo* competitive screening method to identify genes whose inactivation confers a selective advantage to CAR-T cells. Inactivation of 50 genes in a heterogeneous population of T cells expressing an EGFR-targeting CAR revealed that disruption of *REGNASE-1, SOCS1, PTPN2*, and *P16/NK4A* conferred a selective advantage to CAR-T cells in human lung tumor-bearing mice. Consistently, inactivation of these genes improved tumor eradication by CAR-T cells. Interestingly disruption of other genes, described to improve CAR-T cell function in other contexts, had a negative impact in this orthotopic lung tumor model. Further evaluation of long-term effects in a subcutaneous model, highlighted *SOCS1* ablation as the most promising strategy for *in vivo* CAR-T cell amelioration. These results support the importance of evaluating CAR-T cell editing strategies in tumor-specific models and highlight the versatility of our screening approach as a pre-clinical tool for context-specific studies on CAR-T cells amelioration.

## INTRODUCTION

Chimeric Antigen Receptor (CAR) T cells are emerging as an effective treatment for highly refractory and relapsed haematological malignancies, such as leukaemia, lymphomas, and myeloma. Yet, an important fraction of patients still does not respond to these treatments, and despite recent encouraging successes, CAR-T therapy efficacy remains limited in most solid tumors[1]. Clinical trials in leukaemia and lymphomas have shown that a poor expansion and persistence of CAR-T cells in the patient are the main factors predicting a poor prognostic outcome[2]. Coherently, unsuccessful first clinical trials of CAR-T cells against solid tumors report very low numbers of CAR-T cells in patients [3, 4]. On the contrary in different successful clinical trials of CAR-T against solid tumors, [5] especially for anti-GD2 CAR-T cells or anti-IL13Ralpha2 in glioblastoma, CAR-T proliferation and persistence in responding patients were comparable to those of successful anti-CD19 CAR-T therapies, correlating with clinical efficacy[6, 7]. Such results indicate that also in solid tumors, where other obstacles limit their efficacy, a good proliferation and persistence of CAR-T cells remain ultimately essential[4, 8].

Amelioration of CAR-T cells therapy is currently being pursued through several strategies and gene editing approaches are among the most promising.[8–10] Growing number of studies have shown that the ablation of some genes, encoding transcription (co)factors (*/KFZ3, BATF*…)[11, 12], epigenetic modulators (*DNMT3A, TET2*…)[13, 14], signaling regulators (*SOCS1*, DGK-alpha/zeta…)[15, 16], or immunomodulatory receptors (*PD1, T/M3*…)[17, 18], could result in improvement of T or CAR-T cell efficacy, with enhanced expansion or persistence in some *in vitro* or *in vivo* preclinical models. Such results are really promising, but they have been obtained in different experimental conditions, in terms of CAR structure, target antigen, manufacture protocol or tumor model. Since all these elements have been proven to influence CAR-T cell efficacy, it is impossible to directly compare the effects of these different gene candidates[19–23].

Whole genome CRISPR screenings somehow compare the effect of several gene KO simultaneously, such approaches have been successfully applied for in studies on T or CAR-T cells, allowing discovery of new potential candidates target genes[24–30]. Nonetheless, results are not very consistent among these different screenings, suggesting the need for context-specific analyses, in relevant preclinical models. Unfortunately, experiments constraints, such as poor infiltration in tumor tissues or reduced in situ proliferation, pose challenges in *in vivo* solid cancer models[31]. As a result, whole genome approaches would be hardly applicable for specific studies on given CAR-T cells in specific tumor contexts. Notably, large gRNA libraries used in such screenings generally force tailored experimental settings, with expensive and time-consuming sequencing pipelines[27, 28].

Here, we propose a new versatile pre-clinical tool to directly evaluate simultaneously the impact of 50 selected gene modifications in CAR-T cells, through a re-purposing of the CRISPR-KO screening approach. Through this work, we report the application of this strategy to improve CAR-T cells in models of non-small cells lung adenocarcinoma, a promising yet challenging solid tumor target for CAR-T therapy development[32, 33].

## RESULTS

### Set up of a CRISPR-Cas9 system to edit 50 top target genes in a pool of CAR-T cells and a method to quantify subpopulations of edited CAR-T

By an extensive search in the literature in march 2023, we defined a list of 50 genes whose inactivation in T or CAR-T cells could increase their proliferation, persistence or long-term efficiency[11–18, 24–30, 34–66]. (Fig 1 A, Tab. 1) Then, we set up a method to integrate gene editing in the production pipeline of second generation 4-1BB CAR-T cells targeting EGFR. By using a bank of lentiviral vectors expressing a total of 236 gRNAs (4 guides per gene, including 4 negative control genes and 20 non-targeting guides), and transient nucleofection of the Cas9 protein, we produced a pool of CAR-T cells carrying inactivating mutations in one of the selected genes. (Fig. 1 A, B). First, we validated this editing system, successfully targeting *Enhanced Green Fluorescent Protein* gene (*EGFP*) (Fig 1 (). Then, we tested the editing of a gene encoding an epigenetic factor, *R/NF*, for which we confirmed effective editing at a genomic level by Tracking of Indels by Decomposition (TIDE) assay (Fig 1 D).

**Figure 1.**
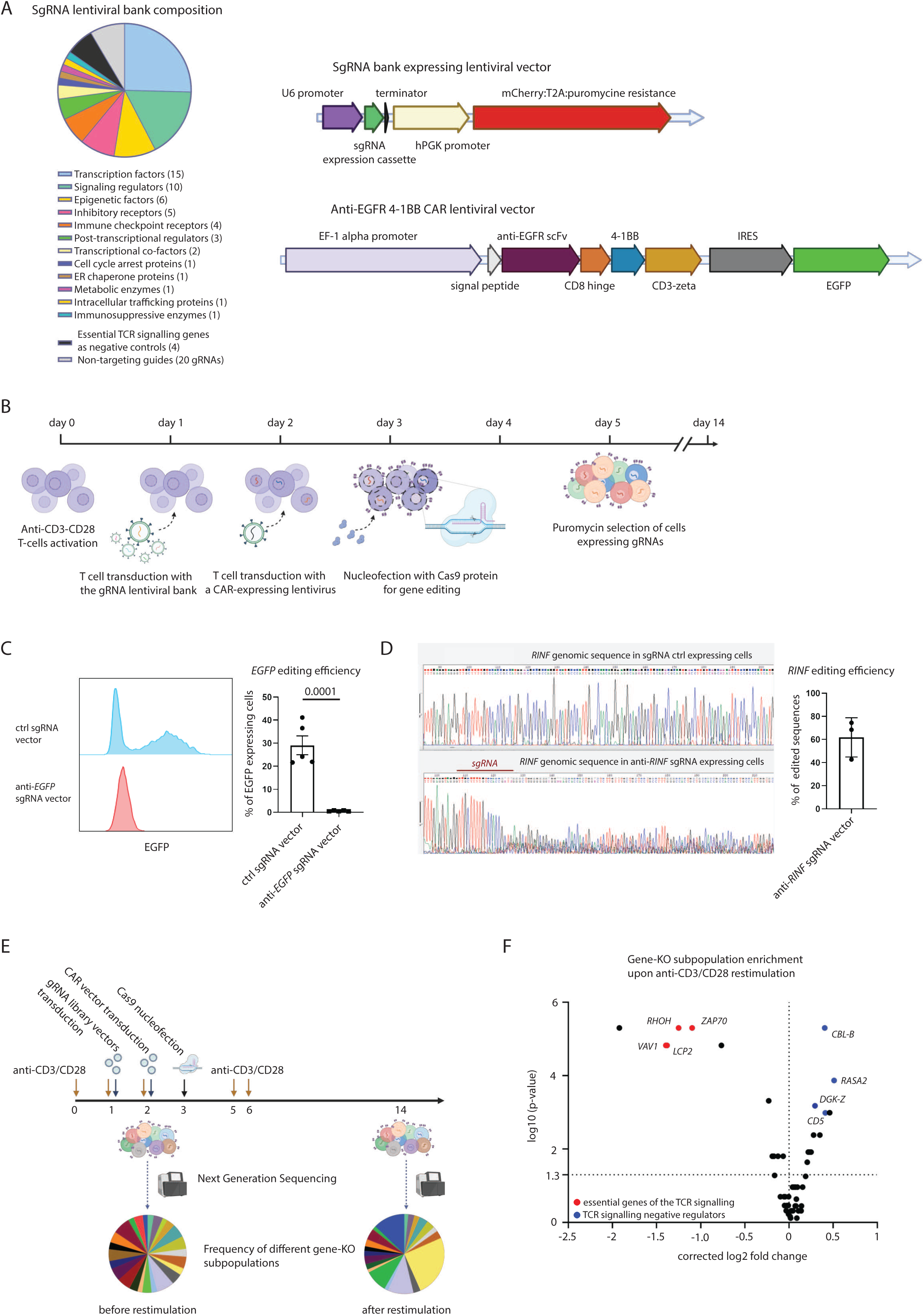
**Description and validation of our CRISPR/Cas9 library**: **(A) (left)** SgRNA bank composition, each of the 50 target genes and the 4 essential negative control genes is targeted by 4 guide RNAs, 20 non-targeting sgRNAs have been added as baseline control. **(right)** Schematics of lentiviral vectors expressing the sgRNA bank and the CAR construct. **(B)** CAR-T cells editing strategy schematics. **(C)** Validation of gene editing strategy by a pilot system targeting *EGFP* in EGFP expressing T cells, as evaluated by signal loss upon knock out trough flow cytometry. p-value of unpaired t-test. **(D)** Validation of gene editing strategy by a pilot system targeting the gene *RINF* in T cells, as evaluated by proportion of modified genomic sequences upon knock out, determined through sanger sequencing and deconvolution analysis of chromatograms files. **(E)** schematics of the in vitro expansion experiment for gRNA library validation **(F)** gene-KO subpopulations frequency variations from library transduction to the end of in vitro expansion assay, genes essential for TCR signalling, used as negative control genes are highlighted in red and TCR signalling negative regulators are in blue. log2 fold change for each gene, expressing frequency changes between the two time points, is corrected to average log2 fold change for 20 non-targeting gRNA. (n=2 biological replicates)

Then, we produced the lentiviral vectors library and set up quantification of CAR-T Knockout (KO) subpopulations by targeted Next Generation Sequencing (NGS). Data were analyzed through Model-based Analysis of Genome-wide CRISPR-Cas9 Knockout (MAGeCK) a bioinformatic tool for gene ranking according to sgRNAs frequencies.[6]

We validated the efficiency of our library by assessing the impact of sgRNA targeting T cell activation genes, previously defined as essential in whole genome CRISPR-KO studies.[24, 26] CAR-T cells inactivated for these genes, *RHOH, LCP2, VAV1* and *ZAP70,* are all significantly negatively selected upon anti-CD3/CD28 driven *in vitro* expansion of CAR-T cells (Fig 1 E,F). Interestingly some candidate genes in this screening are well known negative regulators of T cells activation, and among them KOs of *CBL-B, CD5, DGK-Z, RASA2* were significantly positively selected (Fig 1 F). These results validated our editing system, the lentiviral bank as well as the targeted NGS sequencing strategy and bioinformatic pipeline employed for the analysis.

### Set up and validation of a relevant in vivo orthotopic tumor model of CAR-T therapy

To start, we confirmed that T cells transduced with the 4-1BB anti-EGFR CAR construct efficiently and specifically target EGFR-expressing tumor cells. (Supplementary Fig. 1 A, B) The EGFR antigen is expressed by different solid tumors, including the A549 lung adenocarcinoma cell line. We used this cell line to establish an orthotopic model of lung tumor in immunocompromised NSG mice. (Fig. 2 A, B, (). We observed that our CAR-T cells can infiltrate and accumulate in tumor islets, (Fig. 2 B, Supplementary fig. 1 () and specifically control and eradicate tumors. (Fig. 2 D). We could isolate CAR-T cells from mice lung at the experiment’s endpoint and observed the upregulation of differentiation, activation, and exhaustion markers (Fig. 2 E, Supplementary fig. 1 D). Coherently, such cells show reduced cytotoxicity and proliferation indicating that tumor exposure in this model can induce dysfunction in CAR-T cells (Fig. 2 F).

**Figure 2.**
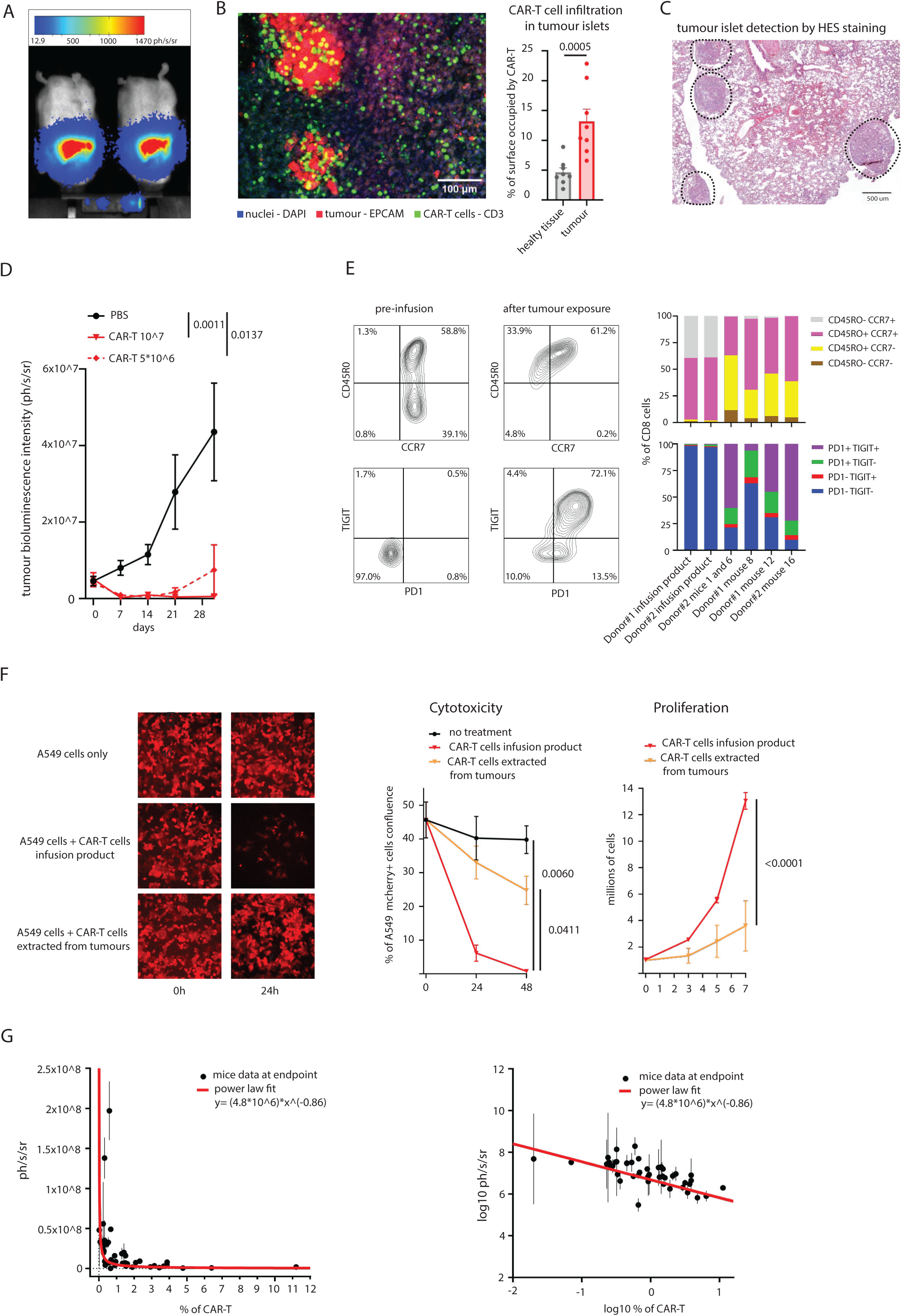
**The A549 orthotopic model is a pertinent model of solid tumour for studying CAR-T cells therapy persistence and efficacy**: **(A)** Exemplary bioluminescence live imaging of A549 cells orthotopic lung tumour. **(B)** Exemplary Immunofluorescence microscopy image of A549 cells (EpCAM staining in red) tumour islet in mice lung, with CAR-T infiltrate (human CD3 staining cells in green). Infiltration analysis has been performed on tumour bearing lung slices images (n=8), 4 days post CAR-T cells injection. P-values are calculated by one-way ANOVA test. **(C)** Exemplary Haematoxylin Eosin staining (HES) of lung tumour section. **(D)** Tumour growth curve, assessed by bioluminescence analysis, in mice receiving mock solution (n=5) or treated with 10˄7 CAR-T cells(n=3) or 5*10˄6 CAR-T cells(n=5). P-values are calculated by two-ways ANOVA test and results for whole curves comparisons are shown. **(E)** Phenotypical characterisation of CAR-T cells by flow cytometry. We compared CAR-T cells from infusion products and CAR-T cells recovered from mice which received the 5*10˄6 dose after long term exposure to tumour *in vivo*. **(F)** Functional characterisation CAR-T cells from infusion products and CAR-T cells recovered from mice. Cytotoxic capacity was assessed by calculating surface reduction of mCherry+ tumour cells when co-cultured with CAR-T cells, at least 3 images were taken per condition, at each time point. Proliferation capacity was assessed by cell counting upon 7 days after anti-CD3/C28 stimulation (n=3 biological replicates for pre-infusion CAR-T cells, n=4 for mice extracted CAR-T cells). P-values are calculated by two-ways ANOVA test, results for final time point comparisons are shown. **(G) (left)** Power-law defined negative correlation among tumour mass, assessed by bioluminescence, and % of CAR-T cells in mice lung at the experiment endpoint. Data shown are collected from all screening experiments performed. The represented power law has been defined through data fitting. (right) representation of the same data and fitting curve on log10 axes.

Interestingly, by gathering data from different experiments, we could observe a strong inverse correlation between the tumour charge and CAR-T cells number in mice lung at the endpoint of the experiment (Fig. 2 G). Such relation appears stronger than an inverse linear correlation, and could instead be described by a power law, as we demonstrated by statistical data fitting (Fig. 2 G). Such observation confirms a strong relationship between CAR-T cell persistence/proliferation and tumor progression in this model.

Overall, these findings support the relevance of this orthotopic lung tumor model for the proposed study.

### The In vivo CRISPR-KO screening assay allows to distinguish1 among 50 top candidate genes1 those whose KO gives the clearest selective in vivo advantage to CAR-T cells

To identify the best target genes to improve CAR T expansion and *in vivo* persistence, we injected tumor-bearing mice with CAR-T cells modified with our lentiviral bank (Fig. 3 A). To increase the power of the test, we produced CAR-T cells from the blood of three healthy donors and we defined sub-optimal therapy conditions, allowing tumor control but not total eradication, for at least five weeks (Fig. 3 B). At the endpoint of the experiments, we could detect substantial proportions of CAR-T in the lungs (Fig. 3 (), which allowed us to quantify each sub-population of the pool of edited CAR-T cells. Briefly, from DNA extracts of the tumor bearing mice lungs, we amplified the sgRNA expressing region of lentiviral vectors in CAR-T cells. PCR amplicons underwent NGS sequencing and data were analyzed with MAGeCK, algorithm.

**Figure 3.**
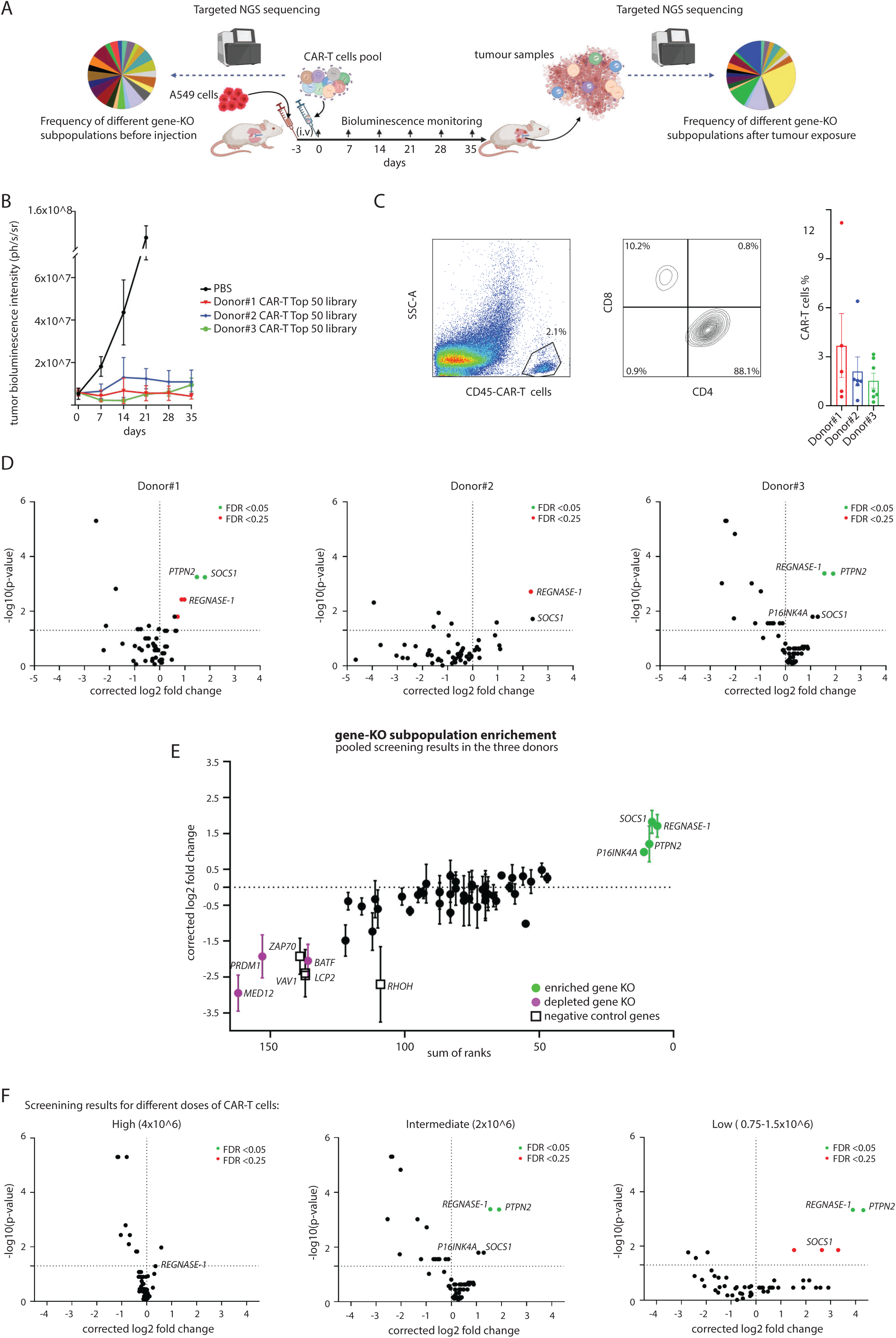
***In vivo* screening in the orthotopic tumour model unveils 4 most promising gene KO strategies**: **(A)** Schematics of the experiment. **(B)** Tumour growth curve as determined by bioluminescence for mice treated either with excipient (n=5) or 2*10˄6 CAR-T modified with our Top 50 library of lentiviral bank, from three different Donors #1 (n=6), #2 (n=6), #3 (n=7) **(C)** CAR-T cells detection in mice lungs by flow cytometry. For CAR-T cells from donor #1, mouse 24 data were collected at anticipated sacrifice at 28 days **(D)** Screening results for CAR-T cells from the three different donors, gene-ko subpopulation proportions are compared in mice lungs at the endpoint of the experiment vs in infusion products. Log2 fold change for each gene, expressing frequency changements between the two time points, is corrected to average log2 fold change for 20 non-targeting gRNA. P-values are determined by MAGeCK permutation test and False Discovery Rate (FDR) values are calculated with Benjamini-Hochberg correction. **(E)** Overall results pooling the three independent screenings, log2 fold change for each gene is calculated as average of these values from each screening. Sum of rank value is determined as sum of the rankings assigned by MAGeCK alpha-RRA algorithm for each gene in the three screenings. **(F)** Screening results from three different experiments in which either 4*10˄6 CAR-T Top 50 library, 2*10˄6 CAR-T Top 50 library or 0.75-1.5*10˄6 CAR-T Top 50 library were injected to mice, with progressively less effective tumour control, and therefore increasing selective pressure on CAR-T cells. Gene-ko subpopulation proportions are compared in mice lungs at the endpoint of the experiment vs in infusion products. Log2 fold change for each gene, expressing frequency changes between the two time points, is corrected to average log2 fold change for 20 non-targeting gRNA. P-values are determined by MAGeCK permutation test and FDR values are calculated with Benjamini-Hochberg correction.

In tumor samples, most of sgRNAs could be detected, even in samples with very low CAR-T cells numbers, proving the good sensitivity of our approach (Supplementary fig. 2 A). We could then define gene targets significantly enriched in different tumor samples of mice receiving CAR-T cells from each blood donor. (Fig. 3 D) Most importantly, we could detect that sgRNA for the KO of the genes *REGNASE-1, SOCS1, PTPN2 and P16/NK4A*, led to a robust selective advantage of CAR-T cells consistently among the three different donors. (Fig. 3 E) As expected we observed depletion of KO of essential genes of the TCR-signaling, used as negative controls. Surprisingly though, some target genes for CAR-T cell amelioration, notably *MED12, PRDM1* and *BATF,* scored as bad or even worse. These genes, while increasing CAR-T cell proliferation or persistence in the experimental conditions of the respective studies,[11, 26, 50] appear rather to impair such capabilities in these specific experimental conditions. (Fig. 3 E).

Throughout the setting up of such screening, different experiments on CAR-T cells from the same donors were performed. We observed that in conditions where CAR-T cells rapidly eradicated tumors, and therefore underwent minimal selective pressure, no meaningful effect could be observed in term of gene-ko subpopulations selection. (Fig. 3 F left, Supplementary fig. 2 B) In more stringent conditions, however, we observed more pronounced effects on gene-ko subpopulations enrichment (Fig. 3 F middle and right, Supplementary fig. 2 B), proving that these outcomes were strictly dependent on selective pressure and are specific to CAR-T cells exposure to tumor.

Overall, these screening results demonstrate the feasibility of our approach and strongly support the value of assessing the effects of gene-editing strategies on specific CAR-T cells and contexts, using models that closely mimic clinical settings.

### Individual disruption of REGNASE-1 and PTPN21 two top hits identified in our in vivo competitive assay1 improved in vivo tumor control capacities of CAR-T cells

To validate whether the selective advantage of top screening hits translated into improved tumor control in vivo, we generated lentiviral vectors expressing gRNAs targeting *PTPN2* and *REGNASE-1*, two of the most reproducible hits across donors and stress conditions. Despite a relatively suboptimal gene editing efficiency, (Supplementary fig. 3 A), both *REGNASE-1* KO and *PTPN2* KO CAR-T cells showed a clearly improved anti-tumor capacity. Compared to unmodified CAR-T cells, both gene-KO strategies led to improved tumor control and extended survival (Fig. 4 A, B) demonstrating the predictive value of our screening approach.

**Figure 4.**
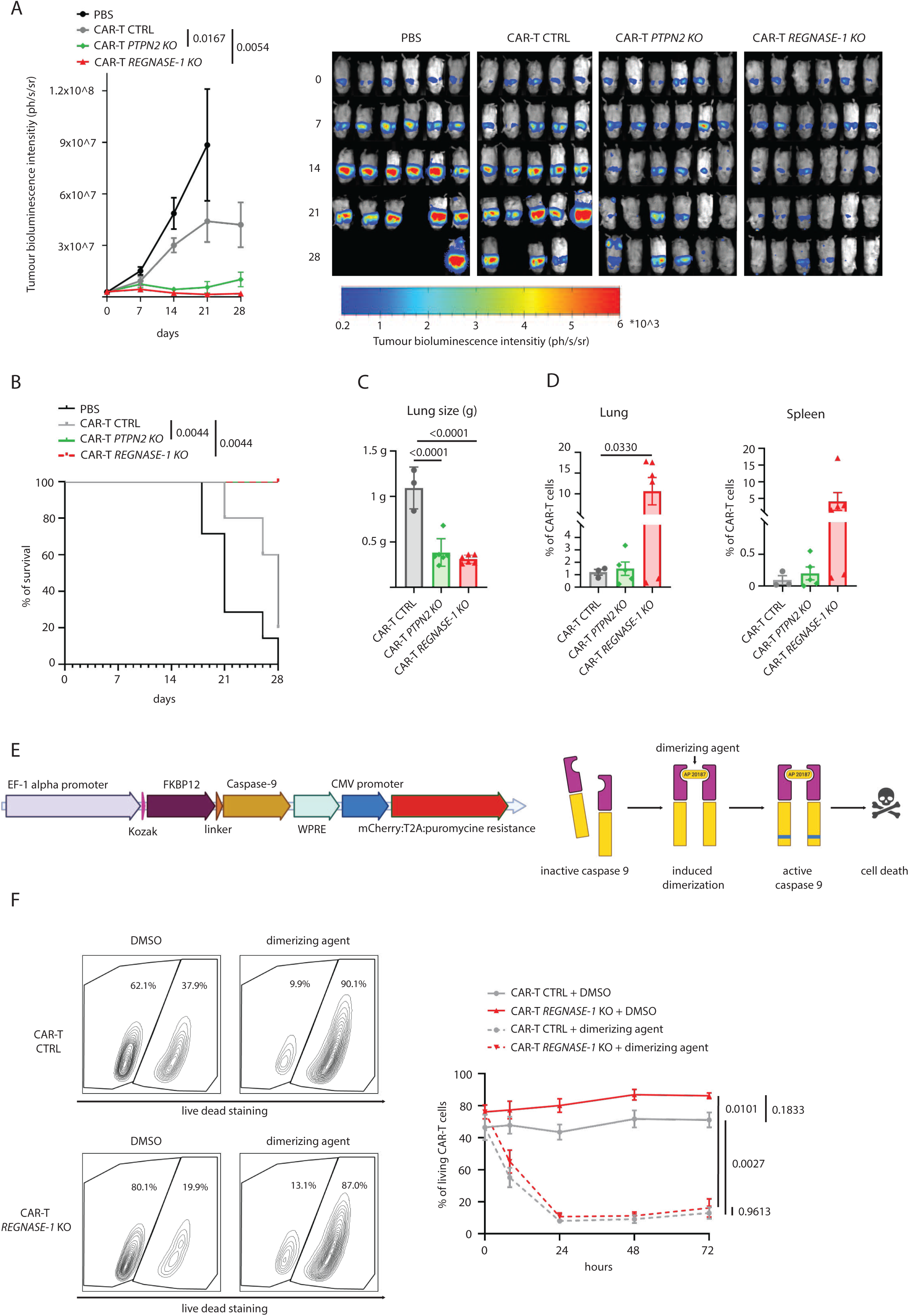
**Validation of top targeting genes *REGNASE-1* and *PTPN2* KO CAR-T cell improved antitumour efficacy**: **(A)** Tumour growth curve in mice treated either with excipient (n=6), 10˄6 CTRL CAR-T cells (n=5), 10˄6 *PTPN2* KO CAR-T cells (n=6) or 10˄6 *REGNASE-1* KO CAR-T cells (n=6). p-values are calculated by two-ways ANOVA test, results for whole curves comparisons are shown. Individual images for each mouse at different time points on the right. **(B)** Survival curves for mice treated with the aforementioned conditions. P-values have been determined by log-rank mantel-cox test. **(C)** Lung size at endpoint, indicative of tumour mass, p-values are calculated by one-way ANOVA test. **(D)** % of CAR-T cells in lungs and spleens of mice at the endpoint as determined by flow cytometry. **(E)** We equipped CAR-T cells with a vector expressing human CASPASE-9 fused with FK506-binding protein 12 (with F36V mutation), to allow dimerization upon administration of the dimerizing agent AP02187, leading to cell death. Unmodified CAR-T cells (n=3) are compared to *REGNASE-1* KO CAR-T cells (n=3). **(F)** Cell death was determined by flow cytometry analysis with live-dead blue dye, data are corrected to cells numbers, to account for dead cell lysis. P-values are calculated by two-ways ANOVA test, results for final time point are shown.

Mice were sacrificed at 28 days to allow comparison with the control condition. We confirmed a markedly reduced tumor burden in the lungs of mice treated with *REGNASE-1* KO or *PTPN2* KO CAR-T cells (Fig. 4 (). Then, we could observe a significantly higher number of *REGNASE-1* KO CAR-T cells at the tumor site and circulating lymphoid organ, as assessed in the spleen (Fig. 4 D), whereas surprisingly *PTPN2* KO had no notable effect on CAR-T cells numbers. These findings suggest distinct mechanisms underlie the improved efficacy conferred by targeting the two genes. *REGNASE-1* KO appears to systematically enhance *in vivo* CAR-T expansion, while *PTPN2* KO may provide a selective advantage under stress conditions without promoting accumulation in in a more favourable context.

The greater number of *REGNASE-1* KO CAR-T cells in treated mice (Fig. 4 D) ensured an effective anti-tumor response, yet it raises the question of potential toxicities.[67] To address this issue, we tested in CAR-T cells combination of *REGNASE-1* KO (Supplementary fig. 3 B), with a safety switch system based on inducible Caspase-9 (Fig. 4 E)[68], that has been successfully applied in clinics.[6] We observed that modified CAR-T retain sensitivity to the iCas9 system (Fig. 4 F), proving these approaches could likely assure safety of modified CAR-T cells therapies.

### REGNASE-1 ablation enhances activation profile in tumor-infiltrating CAR-T cells

To further characterise the effects of *REGNASE-1* and *PTPN2* knockout, we performed Cytometry by Time of Flight (CyTOF) analysis on CAR-T cells isolated from the lungs (tumor site) of treated mice. Uniform Manifold Approximation and Projection (UMAP) was used to summarise data from over 30 markers (Supplementary Tab. 1). Clustering patterns were consistent across samples from each condition (Supplementary fig. 4 A), enabling to focus the analysis on three representative samples and to confirm it in all replicates (Fig. 5 A, Supplementary fig. 4 A). UMAP identified four main clusters: CD4 T cells, CD8 T cells, a minor CD4/CD8 double-positive group, and a very small CD4/CD8 double-negative cluster. The main difference between conditions was a higher proportion of CD4+ cells in *PTPN2* KO CAR-T cells, and more prominently in *REGNASE-1* KO CAR-T cells (Fig. 5 A, Supplementary fig. 4 A). Aside from this, CTRL and *REGNASE-1* KO CAR-T cells more clearly differ, while *PTPN2* KO cells mostly resemble to CTRL cells. (Fig. 5 A, Supplementary fig. 4 A) The first difference in *REGNASE-1* KO CAR-T cells is the emergence of a distinct cluster within the CD4 population. This cluster is defined by co-expression of activation markers, HLA-DR, ICOS, CD28, and immunomodulatory markers CTLA4 and PD1, all upregulated in *REGNASE-1* KO cells (Fig. 5 A, B, Supplementary fig. 4B). Additionally, in these cells we could clearly detect expression of cytotoxic molecules, with some, as granzyme A, showing broader and stronger expression than in other CD4 cells (Fig. 5D). These active, cytotoxic CD4+ cells represent a significantly large fraction of *REGNASE-1* KO CAR-T cells, suggesting a key role in their enhanced expansion and tumor control.

**Figure 5.**
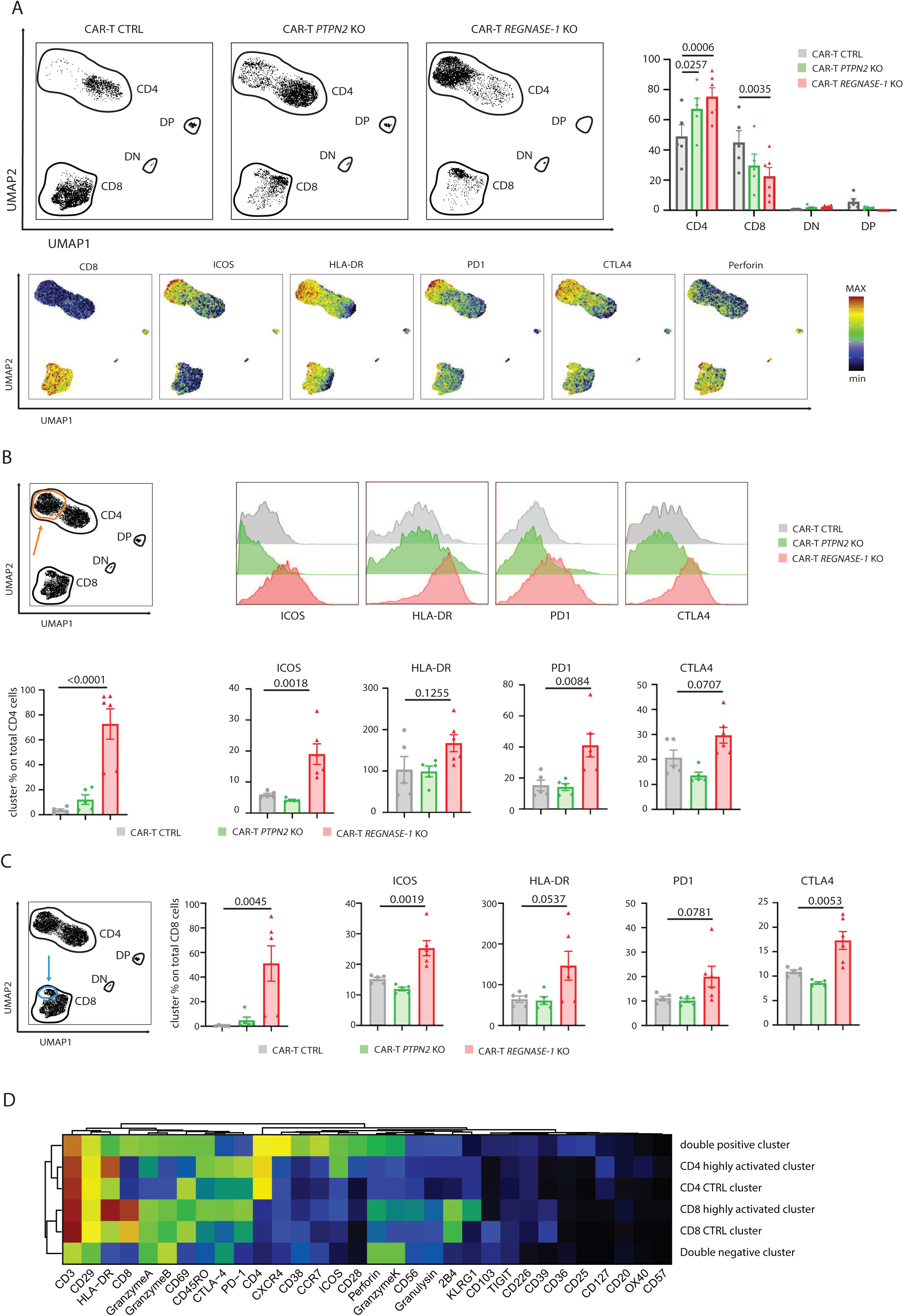
**Cytometry by Time of Flight (CyTOF) characterisation of *REGNASE-1* and *PTPN2* KO CAR-T cells in mice lung tumours**: **(A)** UMAP analysis of CyTOF data from CAR-T cells in lung tumour samples at the endpoint of the experiment. Relative proportion of main clusters ( upper right) and main markers of interests (bottom) in samples from CTRL (n=5), *REGNASE-1* KO (n=6) and *PTPN2* KO (n=5) are also shown. p-values are calculated by one-way ANOVA test. **(B)** Characterisation of variations within the CD4 cluster. the formation of a sub-cluster, for the *REGNASE-1* KO condition, is shown in the combinatory graph of the three exemplary samples (top left) variations of expression for markers defining such cluster in the 3 different exemplary samples are then shown (top right). Frequency of such cluster among all samples and expression levels for the aforementioned markers are shown ( respectively bottom left and bottom right) p-values are calculated by one way ANOVA test. **(C)** Characterisation of variations within the CD8 cluster. the formation of a sub-cluster, for the *REGNASE-1* KO condition, is shown in the combinatory graph of the three exemplary samples (left) overall variations of expression for markers defining such cluster in different condition are then shown (top right). Frequency of such cluster among all samples and expression levels for the aforementioned markers are shown (respectively bottom left and bottom right) p-values are calculated by one way ANOVA test. **(D)** Clustered heatmap and relative dendrogram, data are calculated as overall variation of different markers among defined clusters in all samples.

Within the CD8 cluster, *REGNASE-1* KO cells showed a similar shift, with the emergence of a resembling subpopulation (Fig. 5 (). These cells also show an increase in activation HLA-DR, ICOS, CD28, and immunomodulatory markers CTLA4 and PD1 as their CD4 counterparts and maintained full cytotoxic potential (Fig. 5 (, D). Although representing a smaller fraction of CD8 than CD4 cells, this subset was consistently observed and significantly enriched in *REGNASE-1* KO CAR-T cells compared to CTRL (Fig. 5 (, D, Supplementary fig. 4 (). Despite overall cell number expansion, CD8 T cells made up a smaller proportion of total CAR-T cells in *REGNASE-1* KO samples, suggesting a stronger impact of this modification on CD4 cells. Nonetheless these phenotypic data indicate a notable effect on CD8+ cells as well.

Overall, KO of *REGNASE-1* appears to allow CAR-T cells to achieve and retain a stronger activation upon tumor encounter, which could explain the large increase in CAR-T cells numbers, while maintaining anti-tumor capacities, allowing for better tumor control and ultimately tumor clearance.

### MED12 ablation1 the lowest-scoring gene in our screening1 increases in vitro proliferation and differentiation but reduces in vivo long-term efficacy in this model

Unexpectedly, in our in vivo screen, knockout of at least three promising target genes was as detrimental as inactivation of essential genes used as negative controls (RHOH, ZAP70, VAV1, LCP2). MED12 was the lowest-scoring hit, leading us to test whether its loss could impair CAR-T tumor control in our model. Tumor-bearing mice were treated with a high dose of CAR-T cells, either unmodified or *MED12* KO (Supplementary fig. 5 A). At this dose, unmodified CAR-T cells efficiently cleared tumors within 21 days, with only one mouse relapsing later. Initially, MED12 KO CAR-T cells also reduced or eradicated tumors in some mice. However, tumors later reappeared, persisted, and regrew, leading to death in most MED12 KO-treated mice (Fig. 6 A,B). These results suggest that, under our conditions, *MED12* KO would not impair CAR-T functionality but compromises their long-term persistence in mice.

**Figure 6.**
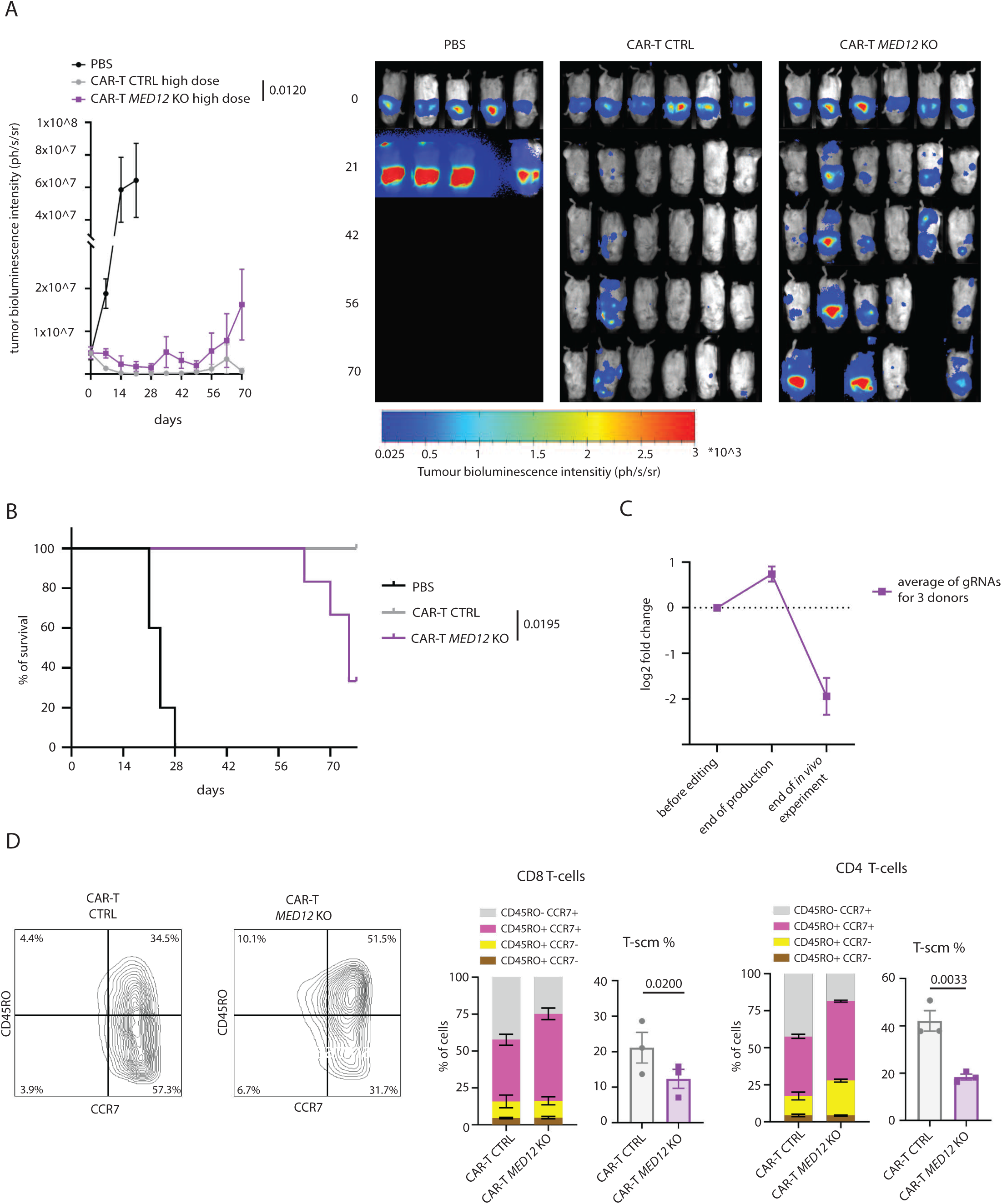
**Study of the effects of *MED12* KO on CAR-T cells *in vitro* proliferation and differentiation and *in vivo* efficacy against lung adenocarcinoma**: **(A)** Tumour growth curve in mice treated either with excipient (n=5), 3*10˄6 CTRL CAR-T cells (n=6) or 3*10˄6 *MED12* KO CAR-T cells (n=6). In reason of the great disparity of signal within mice of the same groups p-value are calculated by non-parametric Wilcoxon signed rank test, results for whole curves comparisons are shown. (nonetheless, p-value of 2-way Anova is also significative, p=0.0002) Individual images for each mouse at different time points on the right. **(B)** Survival curves for mice treated with the aforementioned conditions. P-values have been determined by log-rank mantel-cox test. **(C)** Log2 fold change variations in frequencies of gRNAs targeting *MED12*, through different time points. variations are normalised to non-targeting guides and are calculated as average values for all sgRNA in the three donors. **(D)** Flow cytometry based differentiation profiling for *MED12* KO CAR-T cells (n=3) as compared to CTRL CAR-T cells (n=3) at the endpoint of production phase. T-scm (T stem cell memory) are defined as CD45RA+ and CD27+ t-cells. P-values are calculated with paired ratio t-tests.

To better understand this effect and relate it to prior findings on this genetic modification, we further characterised *MED12* KO CAR-T cells. Firstly, we observed that in the screening, CAR-T cells receiving MED12-targeting gRNAs showed enhanced expansion during production (Fig. 6 (), which we confirmed in an independent experiment (Supplementary fig. 5 B). Although this early in vitro expansion is outweighed by later in vivo depletion, it shows that this modification could be beneficial under different conditions. Phenotypically, MED12 KO CAR-T cells showed greater differentiation, consistent with previous reports [26], with a marked loss of stem cell memory T cells in both CD4 and CD8 compartments (Fig. 6 D, Supplementary fig. 5). This accelerated differentiation may explain the initial proliferation advantage, but also the reduced persistence *in vivo*, as previously shown in the field. [69, 70]

### A high selective pressure in vitro mimics in vivo screening results1 despite attenuated effects

We reasoned that for studying certain characteristics, an in vitro surrogate assay could be informative as long as its results mirrored those from the *in vivo* model. Therefore, we established a high selective pressure *in vitro* assay, restimulating CAR-T cells, by seeding them on irradiated A549 cells, every 5 days over 35 days (Fig. 7 A). Under these conditions, CAR-T cells initially expanded strongly and eradicated tumor cells, followed by stagnation and a clear decline in cell numbers (Fig. 7 B). Analysis of gene-KO subpopulations variations, highlighted results similar to those in vivo: *SOCS1, REGNASE-1, PTPN2* and *P16/NK4A* KO remained top hits, though their advantage was less pronounced. Likewise, genes with negative effects in vivo also scored lower here, though their disadvantage was less consistent between donors (Fig. 7 ().

**Figure 7.**
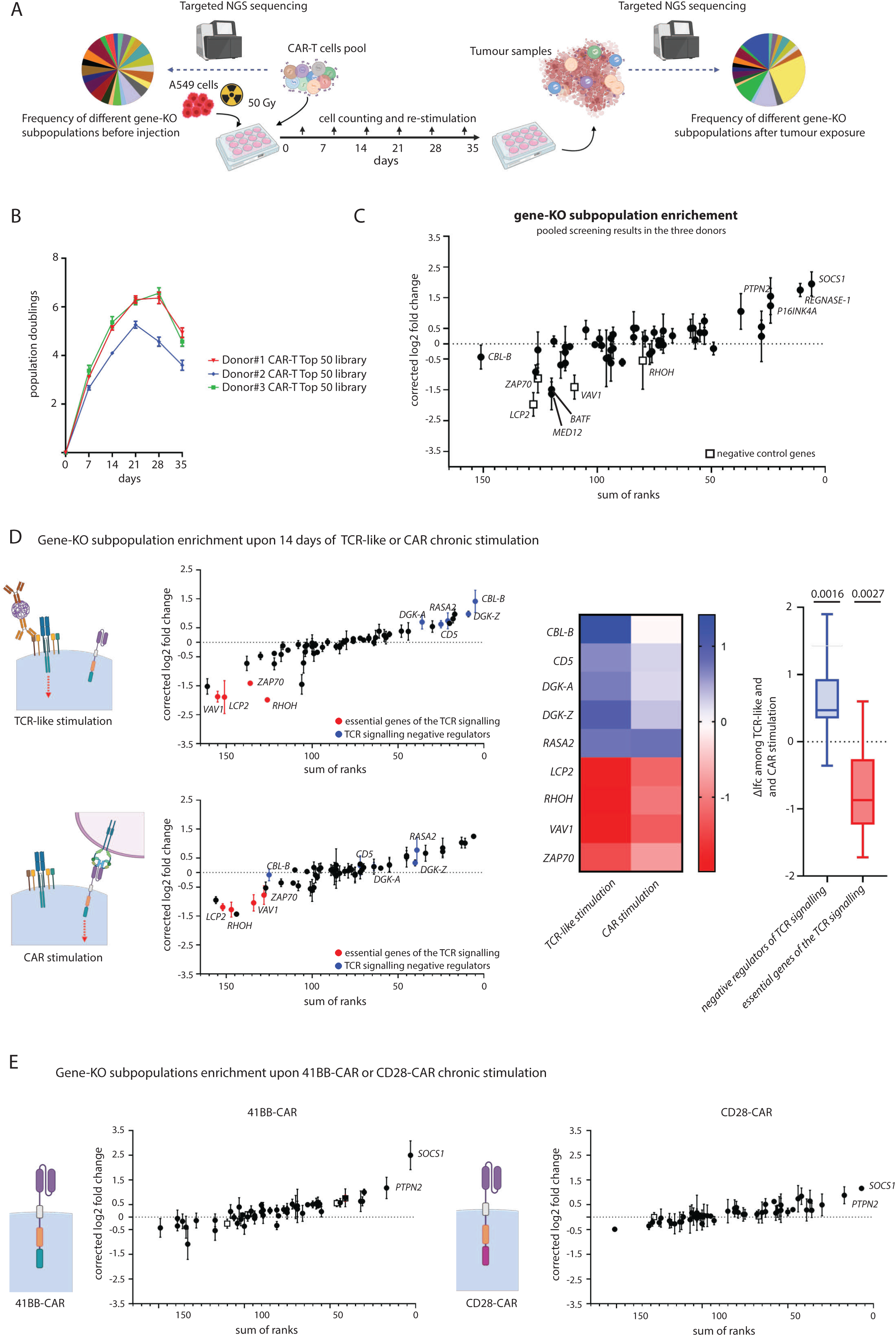
**Application of the screening *in vitro,* for the study of the impact of signalling differences on gene-KO strategies efficacy**: **(A)** Schematics of the experiment **(B)** CAR-T cell proliferation curve as determined by cell counting, 3 replicates for each of 3 donors derived CAR-T cells Donor#1, Donor#2, Donor#3. **(C)** Overall results pooling three independent screenings, log2 fold change for each gene is calculated as average of these values from each screening. Sum of rank value is determined as sum of the rankings assigned by MAGeCK alpha-RRA algorithm for each gene in the three screenings. gene-ko subpopulation frequency variations are compared at the endpoint of the experiment, 35 days, vs in starting products. **(D)** (left) Screening results from three different experiments in which CAR-T cells either were stimulated by anti-CD3/CD28 polymers or irradiated tumor cells, every 3 to 5 days for 14 days, to assess mainly proliferation effects. Gene-ko subpopulation proportions are compared at the endpoint of the experiment vs in starting products. (middle) Heatmap representation of log2 fold change variation for genes either essential to or negatively regulating the TCR pathway. (right) Overall difference in log2 fold change in frequency of CAR-T cells carrying mutation in negative regulators of the TCR pathway and essential genes of this pathway. Variations in CAR-T cells stimulated with anti-CD3/CD28 polymers compared to variations in CAR stimulated CAR-T cells. TCR-like stimulation lead to significantly increased variations for such groups of genes, in both directions. **(E)** Overall results pooling three independent screenings, either with (left) 41BB CAR-T cells or (right) CD28 CAR-T cells. Log2 fold change for each gene is calculated as average of these values from each screening. Sum of rank value is determined as sum of the rankings assigned by MAGeCK alpha-RRA algorithm for each gene in the three screenings. Gene-ko subpopulation frequency variations are compared at the endpoint of the experiment, 35 days, vs in starting products.

### Effects of signaling differences on gene editing strategies efficacy in vitro

After confirming the in vitro assay could sufficiently reproduce the *in vivo* results, we used it to explore how signaling modifications influences gene editing efficacy. First, we asked whether stimulation via the TCR versus the CAR would affect the impact of gene edits on T cell expansion. To mimic TCR signaling, we used anti-CD3/anti-CD28-loaded polymers, while CAR signaling was triggered by exposure to irradiated tumor cells. Notably, after 14 days, knockout of essential TCR pathway genes had a stronger negative effect under TCR-like stimulation than CAR stimulation. Conversely, KO of TCR pathway regulators showed a more beneficial effect in TCR-stimulated cells (Fig. 7 D). Although CAR signaling is designed to mimic TCR engagement, they differ significantly [71, 72], and our findings highlight how such differences influence the efficacy of gene editing strategies, raising a warning on translatability of data obtained from studies on T cells for CAR-T cells amelioration.

We next asked whether these results could be explained by the different co-stimulation signal. Notably, CD28 and 4-1BB co-stimulation is a one of the main differences across CAR-T products [73, 74]. We redesigned our CAR construct to include a CD28 domain instead of 4-1BB and tested these CAR-T cells in the *in vitro* re-stimulation assay. CD28 CAR-T cells showed reduced persistence compared to 4-1BB ones (Supplementary fig. 6 A), which forced to adapt our test, reducing re-stimulation frequency to once every 7 to 10 days (Supplementary fig. 6 B). Under these conditions, gene KO subpopulation changes were less pronounced, only partially reproducing earlier findings. *SOCS1* and *PTPN2* remained top hits, while *REGNASE-1* and *P16/NK4A* KOs showed no clear benefit (Fig. 7 E). Most interestingly though, we observed no major differences in gene-KO subpopulations selected in CD28 and 4-1BB CAR-T cells (Fig. 7 E). These results support previous findings [73, 75] that co-stimulatory domains impact CAR-T persistence, while do not seem to impact effects of gene editing strategies in improving this feature in our settings.

### A subcutaneous tumor model to study gene editing efficacy on longer term1 within different tumor structure and for metastasis control

We sought to translate our screening assay to a subcutaneous A549 tumor model to assess how and if factors like time and tumor architecture affect gene editing strategy efficacy. In this model, CAR-T cells infiltrate tumors sufficiently, though less effectively than in the lung model. Also in this case we found conditions for partial tumor control without full eradication (Fig. 8 A, B). Tumor mass inversely correlated with CAR-T cell numbers, though less strongly than in the orthotopic model (Fig. 8 (). As before, CAR-T cells from tumors showed increased differentiation and exhaustion markers, along with impaired function in ex vivo assays (Supplementary fig. 7 A, B, (). Interestingly, we could observe that subcutaneous A549 tumors developed lung metastases, which were effectively controlled and cleared by CAR-T cell treatment (Fig. 8 D, E).

**Figure 8.**
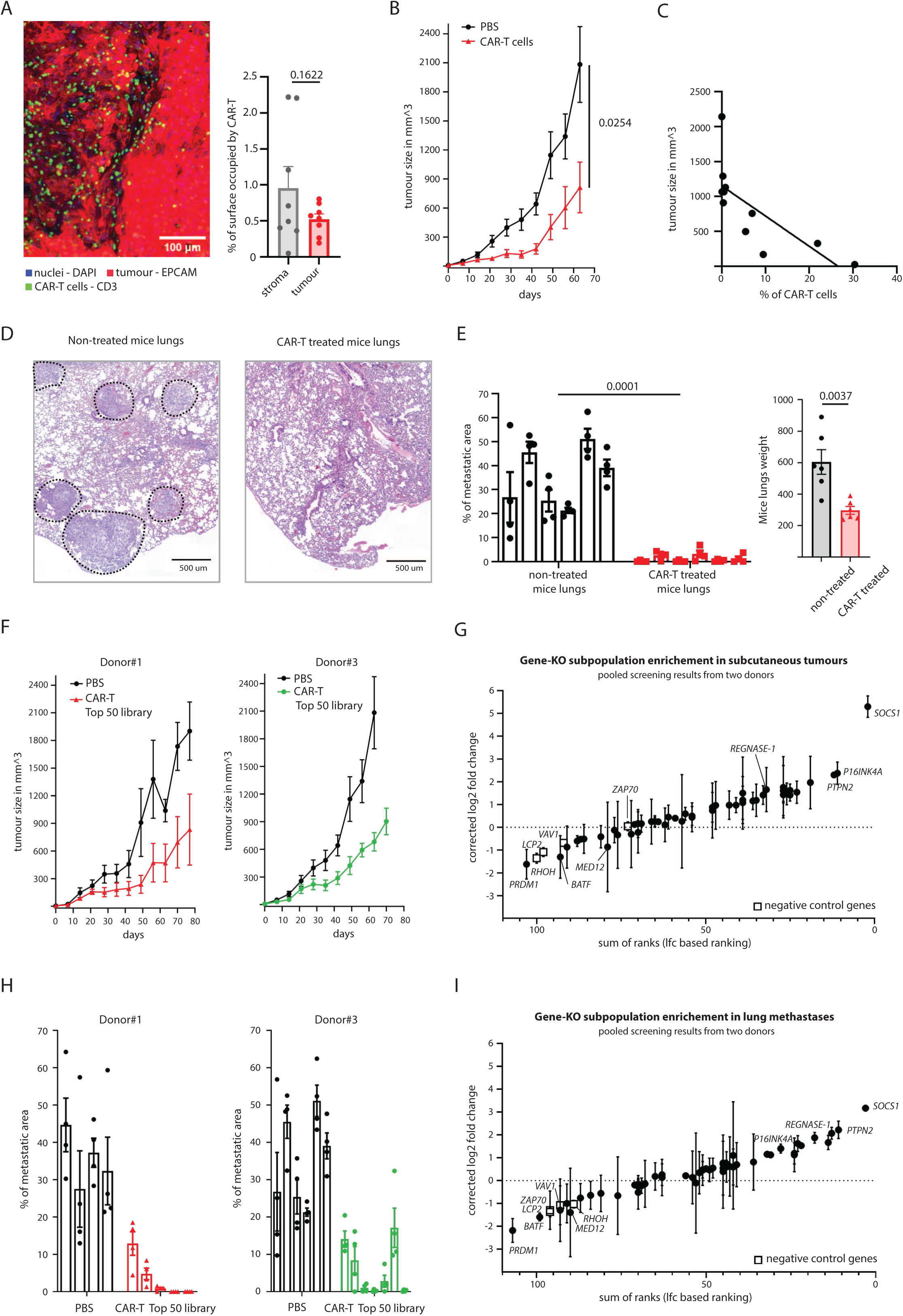
***SOCS1* KO outperforms other gene inactivation strategies in the *In vivo* screening in the subcutaneous tumor model**: **(A)** Immunofluorescence microscopy image of A549 cells subcutaneous tumor, with CAR-T infiltrate and quantification of CAR-T cells infiltration in tumor mass vs stroma (n=8 tumor slices images) **(B)** Tumor growth curve, assessed by caliper measurements, in mice receiving mock solution (n=5) or treated with 7*10˄6 cells(n=5). p-values are calculated by two-ways ANOVA test. **(C)** Negative correlation among tumor mass, and % of CAR-T cells in mice tumor at the experiment endpoint. **(D)** Haematoxylin-Eosin staining of mice lung slices, showing metastatic burden respectively in mice treated with mock or CAR-T cells. **(E)** Machine learning quantification of metastatic area in subcutaneous tumor bearing mice, treated with mock or CAR-T cells. (respectively n=20 and 24 images were used for calculation) p-value is calculated by unpaired t-test. **(F)** Tumor growth curve, assessed by caliper measurements, in mice receiving mock solution (n=5 for both experiments) or treated with modified CAR-T cells 5*10˄6 cells. On the left an experiment with donor Donor#1 CAR-T cells (n=5), on the right the same with donor Donor#3 CAR-T cells (n=6). **(G)** Overall results pooling two independent screenings, log2 fold change for each gene is calculated as average of these values from each screening. Sum of rank is determined as sum of LFC based ranking in each experiment. gene-ko subpopulation frequency variations are compared at the endpoint of the experiment, 70 to 77 days, vs starting products. **(H)** Machine learning based quantification of metastatic area in subcutaneous tumor bearing mice treated or not with CAR-T cells. (n=16 and 20 images for mock and CAR-T treated mice for Donor#1 experiment, n=24 and 24 for the Donor#3 experiment) **(I)** Overall results pooling two independent screenings in the lungs of mice with subcutaneous tumor, log2 fold change for each gene is calculated as average of these values from each screening. Sum of rank value is determined as sum of LFC based ranking in each experiment. Gene-ko subpopulation frequency variations are compared at the endpoint of the experiment, 70 to 77 days, vs starting products.

### Ablation of SOCS1 is the most enriched target gene in the subcutaneous tumour model and for controlling metastases

We repeated our screening in the subcutaneous model, treating mice with modified CAR-T cells and sacrificing them 70–77 days later (Fig. 8 F). Sufficient CAR-T cell infiltration was detected in tumors from 2 out of 3 T cell donors, with high variability in cell numbers (Supplementary fig. 7 D), nonetheless, we could perform sequencing analysis on all these samples. As in the orthotopic model, we confirmed top ranking of *SOCS1, PTPN2* and *P16/NK4*, whereas for *REGNASE-1* KO we could observe a positive effect, but not a consistent ranking across the two donors (Fig. 8 G). The most notable difference was the significantly improved performance of *SOCS1* KO CAR-T cells, suggesting that this genetic modification may offer a more sustained long-term advantage in a complex tumor environment with limited infiltration (Fig. 8 G). Consistent with previous findings, *PRDM1* and *BATF* KOs had a clear negative impact, while *MED12* KO effects were less defined in this context (Fig. 8 G).

We also analyzed the lungs, as we observed the development of lung metastases in this model, detecting CAR-T cells at proportions similar to those in the orthotopic model (Supplementary fig. 7 E) leading to control of the tumor mass (Fig. 8 H). Gene KO subpopulation enrichment analysis confirmed *SOCS1* as the top scorer, followed by *PTPN2* and *REGNASE-1* (Fig. 8 I). These results were consistent with findings in the subcutaneous tumor, making it difficult to distinguish whether selection effects are a result of local proliferation or a consequence of CAR-T cell recirculation from the primary lesion. Interestingly, for genes like *REGNASE-1* and *SOCS1*, the screening results appeared somewhat intermediate between the orthotopic lung tumor (Fig. 3 D) and subcutaneous tumor models (Fig. 8 G).

### Application of the screening assay for the improvement of CAR-T cells mitochondria metabolism1 another relevant phenotype in our context

Finally, we wondered if our library could be applied, in parallel to *in vivo* screenings, to study other CAR-T cell characteristics specifically relevant in a given context, to integrate results from these readouts. We previously found that enhanced mitochondrial respiration improves CAR-T cell migration and tumor control in this model [76]. To investigate this, we set up a screening assay to identify gene KO subpopulations enriched in cells with top 10% high levels of the standard mitochondrial marker MitoTracker expression compared to the bottom 10% (Fig. 9 A, B). Cells in these groups had corresponding differences in mitochondrial content, confirmed by mtDNA quantification (Fig. 9 (). By this screening we identified *PRDM1* and */RF2* as top-scoring genes in CAR-T cells from all three donors (Fig. 9 D).

**Figure 9.**
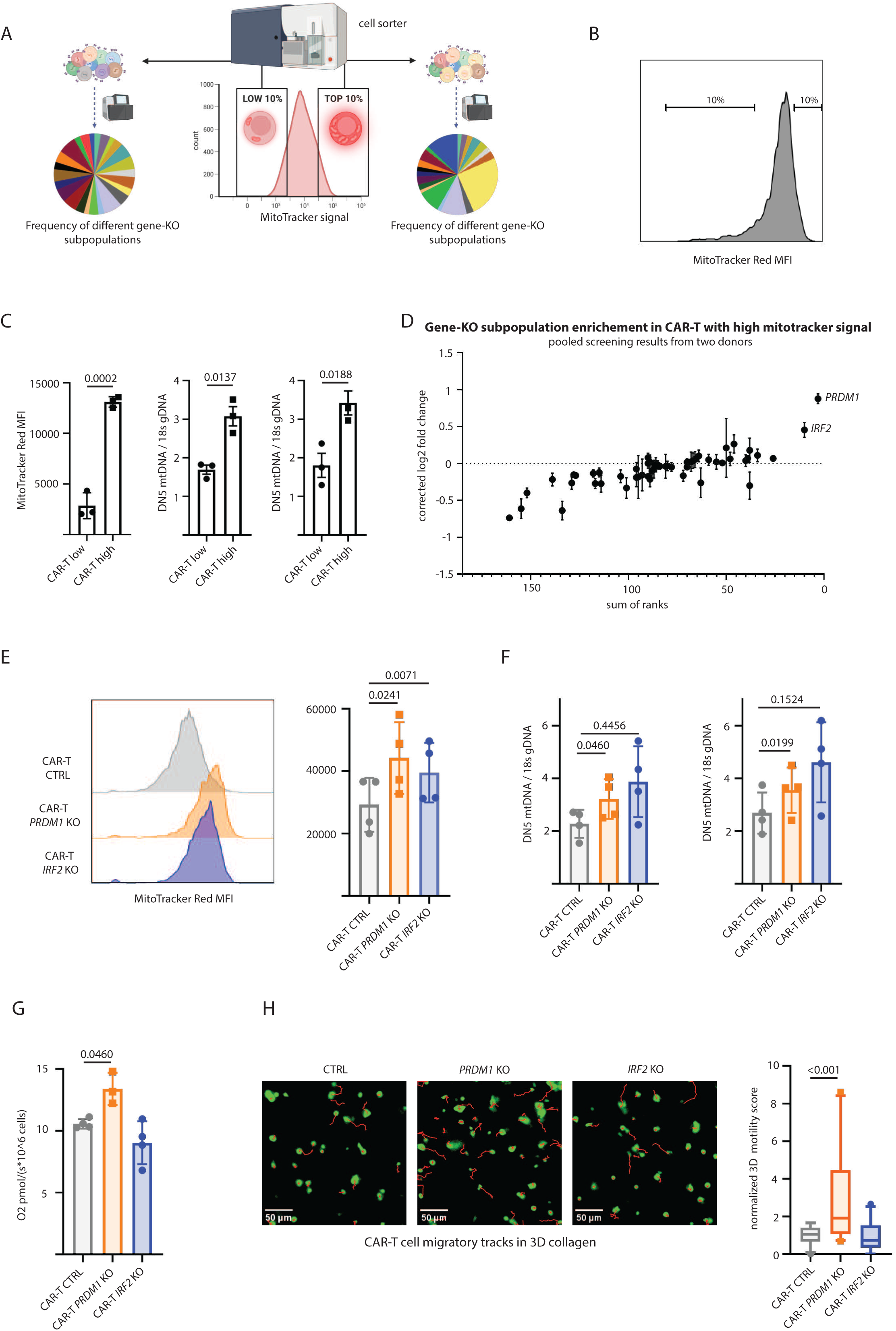
**Screening application to study the improvement of CAR-T cells mitochondrial mass, respiratory and migratory capacity**: **(A)** Schematics of the experiment. **(B)** Exemplary gating for sorting on MitoTracker red signal levels, top 10% and bottom 10% cells have been selected. **(C)** Mitochondrial mass quantification by qPCR measurement of mtDNA copies as compared to genome copies in sorted CAR-T cells. Mitochondrial genes ND1 and ND5 were used for mitochondrial DNA, gene 18s was used for genomic DNA. (n=3 for group) p-values are calculated with one-way Anova test. **(D)** Screening results from three different experiments. gene-ko subpopulation proportions are compared in high vs low MitoTracker signal cells. **(E)** MitoTracker signal levels for CAR-T cells KO for genes *PRDM1* or *IRF2* compared to CTRL ones.(n=4 for group) p-values are calculated with one-way Anova test **(F)** Mitochondrial mass quantification by qPCR measurement of mtDNA copies as compared to genome copies for CAR-T cells KO for genes *PRDM1* or *IRF2* compared to CTRL ones. (n=4 for group) p-values are calculated with one-way Anova test. **(G)** Maximal respiratory capacity as assessed by OROBOROS measurements. (n=3 for *PRDM1* KO, n=4 for CTRL and *IRF2* KO CAR-T) p-values are calculated with one-way Anova test. **(H)** Exemplary migration tracks of CTRL, *PRDM1* KO or *IRF2* KO CAR-T cells in 3D collagen. **(I)** Normalized 3D motility score of CTRL, *PRDM1* KO or *IRF2* KO CAR-T cells. Such score is calculated as product of speed of migrating cell, distance covered and % of actually migrating cells. (n=23 videos) p-values are calculated with one-way Anova on ranks test, with Tukey post-hoc correction.

We then produced individual KO CAR-T cells (Supplementary fig. 8 A) and confirmed increased MitoTracker staining for both *PRDM1* and */RF2* KOs (Fig. 9 E). We also detected higher mtDNA levels in *PRDM1* KO cells, while */RF2* KO showed large variability in mtDNA quantification (Fig. 9 F). Nonetheless, we assessed whether these KOs could enhance CAR-T cell respiratory capacity and *PRDM1* KO cells showed a significant increase in respiration, whereas */RF2* KO had no impact (Fig. 9 G). Finally, we confirmed that the increased respiratory capacity of PRDM1 KO CAR-T cells led to improved migratory capacity in our 3D motility test, while no difference there was no difference for */RF2* KO (Fig. 9 H).

These results support the versatility of our screening for broader studies on CAR-T cells, applying other readouts than just persistence. On the other hand, they highlight the importance of crossing results of these studies, as for persistence readouts *PRDM1* KO proved unsuitable.

## CONCLUSION

The once unforeseeable successes of CAR-T cell therapy come with the need for further improvements, aiming to achieve remission in all leukaemia and lymphoma patients and extend treatment to solid malignancies[1]. The first limitation to tackle, in hematologic malignancies and most importantly in solid cancers, would be the poor proliferation and/or persistence of CAR-T cells in patients[2, 4, 31]. In this context, several groups have made valuable contributions, proposing therapy improvements, with gene editing strategies holding great promise[9, 10].

Alongside discovering new strategies, it will be necessary to evaluate them in specific contexts. Indeed CAR-T therapies widely differ in terms of the tumor and antigen targeted, the CAR construct design, the production pipeline, and predicted patient profiles, all impacting their efficacy. [19–23] Different whole genome CRISPR-KO approaches have been applied to study T or CAR-T cell improvements, and key lessons can be drawn from them. First, that applying such approaches *in vivo* requires ingenious strategies to overcome limitations intrinsic to these models [25, 27–29, 77]. Second, that context specific studies would be crucial, as results from these investigations do not, or only partially confirm one another. [24–29, 77] In this work, we propose a method for focused small-scale studies in pertinent models to identify genetic modifications effective for a given CAR-T therapy. This approach allowed us to identify genes *REGNASE-1, SOCS1, PTPN2*, and *P16/NK4A* that gave CAR-T cell a long-term selective advantage in an orthotopic tumor model. Later, we performed proof-of-concept validation of two out of these genes, *REGNASE-1* and *PTPN2* confirming that their individual could enhance CAR-T cell antitumor efficacy. Suggesting that genes identified by our screening would be prime targets for improving CAR-T therapy in lung adenocarcinoma.

Surprisingly, we revealed that some genes, such as *MED12*, *PRDM1*, and *BATF*, led to negative selection in our screening. While these results do not undermine the potential of these modifications, given the convincing positive effects on CAR-T cells in the context of other studies, [11, 26, 54], they emphasize the need for context-specific analyses.

An approach like the one proposed can also provide insights into the underlying biological mechanisms of CAR-T success or failure. For instance, we observed that *REGNASE-1* KO CAR-T cells had a marked over-representation of highly activated, pre-exhausted, and cytotoxic CD4 and CD8 cells in the tumor site, correlating with antitumor efficacy in the orthotopic lung tumor model. This increased activation could explain the improved proliferation of these cells, and does not seem to compromise cytotoxic properties, ensuring antitumor efficacy. The upregulation of immunomodulatory markers seems a consequence of high activation, not detrimental to these qualities. These cells share characteristics of CD8 Tpex cells [78], associated with improved persistence of CAR-T cells and better tumor control, reportedly enriched in *REGNASE-1* KO CAR-T cells in mice [79].

Conversely, MED12 KO CAR-T cells showed accelerated differentiation during the production phase, which may explain their early in vitro proliferation and potentially the improved efficacy in other models [26], but proved deleterious in our orthotopic lung tumor model.

Another key aspect for CAR-T therapy development is the need to use diverse models and assays, each addressing specific aspects of the treatment[80, 81]. Therefore, CAR-T cells improvement driven by gene-modification strategies could benefit from integrating multiple readouts.

For CAR-T therapy in a given pathology, different *in vivo* models may be valuable. The orthotopic lung tumor better mimics CAR-T trafficking in lung cancer and tumor growth in a more physiological niche.

[82] Conversely, subcutaneous models suit longer-term studies and better reflect structural barriers hindering CAR-T therapy in solid tumors. [83, 84]

In this study on lung adenocarcinoma, we used both models and found that while *SOCS1* KO matched *REGNASE-1* KO in the orthotopic model, it significantly outperformed it in the subcutaneous one. While it is hard to predict which model would better reflect real life conditions in patients, combining both readouts improves confidence in identifying *SOCS1* KO as a promising strategy for CAR-T enhancement in lung adenocarcinoma.

One major promise of CAR-T therapy in solid tumors is widespread tumor eradication and enhanced immunosurveillance, eliminating metastases along with primary lesions to ensure relapse-free remission [85, 86].

In this study, we assessed CAR-T cell genetic modifications enriched in both primary tumor and metastatic site within the same mice. We observed a general concordance of results, possibly due to local selection of similar genes or CAR-T cell recirculation. Interestingly, for KO of *SOCS1* and *REGNASE-1*, results in lung metastases were intermediate between those in the subcutaneous tumors and the lungs in the orthotopic models. This may reflect niche-specific preferences for certain gene editing strategies and further studies in this direction would be of great interest.

Further addressing the issue of context-specific readouts, various examples arise: in some settings, PD-L1-induced inhibition may be the main obstacle, while in others it may be nutrient deprivation or hypoxia within the tumor site [87–89]. Developing gene editing strategies to tackle these challenges could be valuable as secondary readouts to *in vivo* persistence.

In our tumor model, we previously demonstrated the importance of strong mitochondrial metabolism and CAR-T cell migration[76]. Using a screening based on high MitoTracker signal, we identified *PRDM1* KO as enhancing mitochondrial mass, respiration, and migration. However, our other screenings showed this modification impairs CAR-T cell expansion and persistence, making it unsuitable for improvement in this context.

Results from different readouts may also guide more complex editing approaches, combining multiple gene KOs. It was shown that KO of *REGNASE-1, PTPN2*, and *SOCS1*, prioritized in our study, can be combined with additive effects[28]. Given current limitations of CAR-T therapy in lung tumors, such a’radical’ approach could be envisaged. In this work, we also demonstrate the efficacy of a suicide gene to eliminate both REGNASE-1-modified and non-modified CAR-T cells. This safety switch could be essential to support development of these highly effective CAR-T cells while ensuring toxicity control.

To conclude, we propose a new CRISPR-KO screening approach in CAR-T cells, tailored to improve specific therapies in relevant *in vivo* models. Notably, a recent study (preprint [90]) using a similar strategy in multiple myeloma also confirmed the importance of *REGNASE-1*, *SOCS1*, and *PTPN2*, while *MED12* inactivation showed neither a positive nor negative effect, despite inclusion in their gRNA library. These data suggest some gene KOs may have broad efficacy, even though we could observe quantitative differences of such effects in different models. Others may be highly context-dependent, ineffective or even harmful in some settings, but beneficial in others, highlighting the value of context-specific studies.

## MATERIALS AND METHODS

### Study approvals

All human studies complied with French biomedical research regulations and the principles of the 1975 Helsinki Declaration and its revisions. Peripheral blood samples from adult donors were obtained from the Etablissement Frani;ais du Sang (EFS) under agreement 18/EFS/030, with informed consent for research. Samples were anonymized.

NOD.Cg-PrkdcA(scid)Il2rgA(tm1Wjll)/SzJ (NSG) mice were purchased from The Jackson Laboratory (strain 005557) and housed under pathogen-free conditions at the Cochin Institute Animal Facility, in same-sex cages of 5–6 mice. Environmental parameters were regulated per FELASA guidelines, with cage enrichment provided. Health monitoring and pathogen screening were routinely conducted. Animal discomfort was minimized, and usage was reduced in line with Directive 86/609/EEC. Both sexes were used, with balanced distribution among experimental groups, and euthanasia was performed by cervical dislocation. The protocol was approved by Paris Descartes University (CEEA 17-039) and the French Ministry of Research (APAFiS 15076 and 49768).

### Cell Culture and Reagents

Peripheral blood mononuclear cells (PBMCs) were isolated via Ficoll gradient centrifugation (800 g for 20 min, RT) and cryopreserved in 90% fetal bovine serum (FBS) (Thermo Fisher, A3840002) and 10% DMSO (Thermo Fisher, 327182500). For CAR-T cell production, PBMCs were cultured in RPMI 1640 medium (Thermo Fisher, 21875) with 10% FBS, 2 mM L-glutamine (Thermo Fisher, 25030081), and 100 U/ml penicillin/streptomycin (Thermo Fisher, 15140130), supplemented with 25 ng/ml IL-7 (Miltenyi, 130-095-363) and IL-15 (Miltenyi, 130-095-765). Cells were activated 24 hours post-thaw with T cell TransAct (Miltenyi, 130-111-160) at a 1:200 dilution for 48 to 96 hours.

For experiments with the sgRNA-expressing lentiviral library or the iCasp9-expressing virus, 1 ug/ml puromycin (Thermo Fisher, A11138-03) was added two days post-nucleofection for three days, then reduced to 0.5 µg/ml until the end of CAR-T cells production. Human A549 (mCherry⁺, Luciferase⁺) and HEK-293T tumor cells were cultured in DMEM medium (Thermo Fisher, 11995) with 10% FBS, 2 mM L-glutamine and 100 U/ml penicillin/streptomycin. All cell lines were tested for mycoplasma contamination.

### T cells viral transduction and gene editing

For experiments with the sgRNA-expressing lentiviral library or individual sgRNA-expressing viruses, cells were activated, then infected at 24h with 2.5 µl/ml of lentiviral pool or individual sgRNA lentiviruses. At 48h, the medium was changed, and cells were infected with 100 µl/ml of CAR-expressing viruses and lentiBOOST (Revvity, SB-A-LF-901-01) at 1:200. At 72h cells were washed, divided into 5 × 10u aliquots, resuspended in 16.4 µl P3 and 3.6 µl Supplement Reaction solutions from P3 Primary Cell 4D X Kit S (Lonza, V4XP-3032), and 5 µg of Cas9 protein (Thermo Fisher, A36497). Electroporation was performed using the Amaxa 4D Nucleofector (Lonza), following the manufacturer’s instructions. After 30 minutes of recovery at 37°C, cells were cultured for 24 more hours under stimulation conditions. For procedures without gene editing, only CAR virus infection was performed. In the suicide gene experiment, cells were infected at 24h with 100 µl/ml of iCASP9-expressing viruses, at 48h with CAR-expressing viruses, and at 72h sgRNAs (Thermo Fisher, A35534) were added in a 3:1 molar ratio to Cas9 protein during nucleofection. The REGNASE-1 sgRNA used for RNP editing had target sequences 5’-AAGGAGGTCTTCTCCTGCCG-3’ and 5’-CAGTGTTTGTHCCATCCTGG-3’.[67]

### lentiviruses design and production

To generate anti-EGFR 4-1BB-CAR, scFv sequences from nimotuzumab Ab (IMGT/2Dstructure-DB INN 8545H, 8545L) were introduced into the expression cassette encoding CD8 hinge and transmembrane domains (194–248 Aa, GenBank: AAH25715.1), 4-1BB costimulatory domain (214– 255 Aa, GenBank: AAX42660.1), CD3z signaling domain (52–163 Aa, GenBank: NP_000725.1), and GFP coding sequence (downstream of IRES).

To generate anti-EGFR CD28-CAR, 4-1BB was replaced with CD28 (GenBank: NM_001243078.2) in an identical new vector (Vector Builder VB240303-1098rav).

Lentivirus expressing FK506-binding protein 12 (F36V) fused with human CASPASE-9 protein (1-134 AA deleted) and linked to HA tag was purchased as commercially available (Vector Builder VB900176-7377zdg).

Lentiviral bank expressing gRNAs (Vector Builder Lib230420-1523hhp) was custom designed by Vector Builder, targeting 50 genes of interest and 4 essential genes in T-cell activation (LCP2, RHOH, VAV1, ZAP70). For each gene, 4 gRNAs were designed, including 2 previously described in the literature when available. 20 non-targeting gRNAs were also included as controls.

Custom sgRNA expressing plasmids were designed on demand, including sgRNA targeting *EGFP* (Vector Builder VB230314-1357xnw, sequence: 5’-GGGCGAGGAGCTGTTCACCG-3’), *R/NF* (Vector Builder VB230314-1123mnq, sequence: 5’-GTTGCTTTTGTCCACCGCCA-3’), scramble (Vector Builder VB240125-1049dfy, sequence: 5’-ACGTGGGGACATATACGTGT-3’), *REGNASE*-1 (Vector Builder VB240125-1038zad, sequence: 5’-TTCACACCATCACGACGCGT-3’), *PTPN2* (Vector Builder VB240216-1035aey, sequence: 5’-AGTGCAGGCATTGGGCGCTC-3’), *MED12* (Vector Builder VB240715-1614nfg, sequence: 5’-TAACCAGCCTGCTGTCTCTG-3’), *PRDM1* (Vector Builder VB240918-1470wdv, sequence: 5’-TCGATGACTTTAGAAGACGT-3’), and */RF2* (Vector Builder VB240918-1473bwg, sequence: 5’-GGATGCATGCGGCTAGACAT-3’).

Lentiviral particles were produced by transient co-transfection (polyethylenimine:DNA at 3.5:1 ratio) of HEK 293T cells with the plasmids expressing the sequence of interest and second-generation packaging plasmids psPAX2 and pMD2.G (Addgene, 12260 and 12259). Viral supernatants were collected at 48 and 72 h post-transfection, concentrated by ultracentrifugation at 25,000 g for 2 h, and stored at 280 °C.

### Gene editing assessment by deconvolution assays

DNA extraction from bulk-edited and control CAR-T cells was performed using the FastPure Blood/Cell/Tissue/Bacteria DNA Isolation Mini Kit (Vazyme DC112-01-AB). Cells were lysed with proteinase-K (Vazyme, DE102-01) at 56°C, and DNA was isolated using silica gel column purification. Gene editing efficiency was assessed by PCR amplifying the genomic region surrounding the CRISPR-Cas9 sgRNA target site; then PCR amplicons were then subjected to Sanger sequencing (Eurofins tubeSeq supreme service). Sequencing data were analyzed using DECODR (ChristianaCare) [91] via the Latch.bio online console. DECODR uses a deconvolution algorithm to interpret mixed sanger sequencing chromatograms, quantifying indel frequencies.

### Orthotopic and subcutaneous tumor models

For the orthotopic tumor model, 4 × 10˄6 human A549 cells (mCherry+ Luciferase+) were injected intravenously (i.v.) into the tail vein of 6 to 10 weeks-old NSG nude mice (balanced sex distribution across conditions) in most experiments. In a screening experiment, 20 x 10A6 A549 cells were injected instead.

For tumor growth follow-up, 3 days after i.v. injection of A549 cells, mice were injected i.v. with anti-EGFR CAR T cells (number specified in the text). Tumor growth in the lungs was monitored by bioluminescence using the PhotonIMAGER system (Biospacelab) and M3Vision software (Biospacelab, version 1.2.1.32016). Tumor burden was evaluated using a grid based on bioluminescence, behaviour, appearance, weight, etc., with mice sacrificed upon reaching the endpoint threshold score.

For the subcutaneous tumor model, 5 × 10˄6 A549 cells were injected subcutaneously (s.c.) into the right flank of 2-month-old NSG nude mice (balanced sex distribution). For infiltration analysis, after 5/6 weeks, mice were injected i.v. with 5 × 10˄6 anti-EGFR CAR T cells and sacrificed 4 days later for the same analyses as the orthotopic model. For tumor growth follow up, 14 days post-injection, A549 tumor-bearing mice were i.v. inoculated with 7 × 10˄6 anti-EGFR CAR T cells or 5 × 10˄6 CAR T cells modified with the lentiviral bank, and tumor growth was followed by caliper measurement for 9-11 weeks. An evaluation grid was also used for this model, adapted for subcutaneous tumor mass measurement, with mice sacrificed upon reaching the endpoint score.

### Infiltration analysis

For tumor infiltration assessment 3 weeks after tumour injection (4 × 10˄6 human A549 cells (mCherry+ Luciferase+)), mice were infused i.v. with 5 × 10˄6 anti-EGFR CAR T cells, and after 4 days, they were sacrificed and lungs collected.

Lung or subcutaneous tumor samples were fixed overnight (o.n.) with PLP solution (1% paraformaldehyde, 2 mM lysine-HCl, 550 mg/L NaIO4) at 4 °C, cut into 1–2 mm³ pieces, embedded in 7% agarose (Sigma Aldrich A0701), and sectioned into 150 µm-thick slices using a vibratome (VT 1000 S, Leica). For extracellular staining, slices were incubated for 1 h at RT with 0.5% BSA in DPBS. The slices were placed onto Millicell 0.4 µm organotypic inserts (Millipore PICMORG50), and 30 µl of 0.5% BSA/DPBS with the desired Abs were added. A metal ring kept the solution in place, and Abs were incubated o.n. at 4 °C. The following day, slices were washed twice in DPBS and mounted on Superfrost microscope slides (Epredia J1800AMNZ) using Vectashield antifade mounting medium (VectorLabs H-1900). The following antibodies were used (200 ng/slice): FITC-anti-CD3 (Biolegend 344804, clone SK7) and PE-anti-EpCAM (Biolegend 324206, clone 9C4). 4′,6-Diamidino-2-phenylindole (1/1000) was used to stain nuclei. Images were captured with a spinning-disk (Yokogawa CSU-W1 T1) Ixplore IX83 microscope (Olympus) equipped with an ORCA-Flash 4.0 V3 (sCMOS Hamamatsu) camera and CellSens Dimension software (Olympus, v4.1.1). CAR-T cells infiltration was analyzed using an ImageJ threshold mask to identify EpCAM+ areas (tumor islets) and CD3+ areas (T cells). <Subtract= or <multiply= ImageJ commands were used to define T cells in tumor stroma or islets, respectively. Raw data from these infiltration experiments were previously employed as control condition in figure 7 B and E of another publication from our group [76], Data have been here used for a different analysis to study and compare CAR-T cells infiltration within the tumour tissue in the two models.

### HES staining on tissue samples and machine learning metastatic burden quantification

Tissues were fixed in 4% PFA for 24 hours, dehydrated in graded ethanol, isopropanol, and xylene (Thermo Fisher, A11358.0F), then embedded in paraffin for FFPE storage. Sections were cut into 6 slices (4 µm thick, 150 µm apart), using a microtome (Autosection, Sakura). Slides were stained on the SPECTRA stainer (Leica) with H&E, deparaffinized, rehydrated, rinsed, incubated in Mayer’s haematoxylin for 3 minutes, rinsed, stained in erythrosine for 5 minutes, followed by 5 minutes in 1% alcoholic saffron. Digitalized Images were obtained Stained slides were scanned at 20x magnification using the Lamina Slide Scanner (Perkin Elmer) and analyzed with Case Viewer software (v 2.2).

From these images, 4 cropped areas (4.5x magnifications) from at least 2 different slices were analyzed with InForm® software. Cropped images were converted to optical density with eosin and purple haematoxylin channels. Tissue segmentation was performed by categorizing regions as no tissue, healthy tissue, and tumor tissue. At least 10 areas per category were used to train the machine learning algorithm. Segmentation was then automated and visually inspected, with corrections made manually. The percentage of metastatic area was calculated as the ratio of metastatic area to the sum of healthy and tumor tissue areas.

### Tumors and organs dissociation

Tumor-bearing mice lungs were cut into 1-2 mm pieces, digested with Liberase (Roche 5401119001) at 37 °C for 30 minutes, then smashed using a 70 µm Cell Strainer filter (Corning 352350). Collected cells were stained for flow cytometry (BD Fortessa) or pelleted for DNA extraction. Subcutaneous tumors were processed similarly to obtain single-cell suspensions, stained or pelleted for DNA extraction. Spleens were collected, smashed using a 70 µm filter, and cells were processed as above.

### Tumor infiltrating CAR-T cells isolation

CAR-T cells were isolated from single-cell suspensions using the human CD45 MicroBead Kit (Miltenyi, 130-121-563) and LS Columns (Miltenyi, 130-042-401). Cells were resuspended in MACS buffer, incubated with CD45.1 MicroBeads at 4°C for 15 minutes applied to LS Columns, washed, and eluted with buffer. Collected CAR-T cells were either stained for flow cytometry (BD Fortessa) for the lung tumor characterisation experiment or cultured for 48 hours and used in cytotoxicity and proliferation assays for both lung and subcutaneous tumor experiments.

### Cytotoxicity assays

Cytotoxic activity of in vitro expanded and tumor-extracted anti-EGFR 4-1BB-CAR T cells was assessed using mCherry-expressing A549 cells. A549 cells were seeded at 20 × 10u cells per 250 µl in 96-well plates (Corning 351172) with DMEM. After 48 hours (time 0 h), plates were irradiated with 50 Gy using a BIOBEAM GM 2000 (Ekert and Ziegler). anti-EGFR 4-1BB-CAR T cells were added at a CAR:target ratio of 4:1 in RPMI medium, without cytokines. In the experiment involving CAR-T cells extracted from subcutaneous tumors, ratios of 1:2 were used, because of the low number of CAR-T cells recovered. A549 cell viability was determined by measuring the mCherry-positive area at different time points: A fluorescence microscope was used in the experiment shown in Figure 2, and an Incucyte S3 (Sartorius) was used for all others. The mCherry-positive area was compared between wells with and without CAR T cells.

### In vitro proliferation assays

In vitro cultured and tumor-isolated anti-EGFR 4-1BB-CAR T cell proliferation was evaluated by trypan blue corrected living cell counting. Cells were cultured for 48 hours in complete medium, then stimulated with anti-CD3/CD28 conjugated TransAct (Miltenyi, 130-111-160) at 1:200 for 48 hours. Proliferation was followed for 7 days after activation.

### Flow cytometry

FSC-A/SSC-A gating and LIVE/DEAD™ Fixable Blue Dead Cell Stain Kit (Invitrogen, L23105) were used to define live and dead cells. Extracellular proteins were stained with antibodies (1:100 dilution): BV785 CD45 (Biolegend, 304048), PE CD45 (Biolegend, 368509), BV650 CD4 (Biolegend, 317436), BUV737 CD8 (BD Biosciences, 612754), BV711 CD45RA (Biolegend, 304138), APC CD45RO (Biolegend, 304210), APC-Cy7 CD27 (Biolegend, 302816), PE-Cy7 CCR7 (Biolegend, 353225), APC PD-1 (Miltenyi, REA1165), BV421 TIM3 (Biolegend, 345007), BV786 LAG3 (Biolegend, 369321). T cell subpopulations were defined as follows: stem cell memory-like (Tscm, CD45ROnegCCR7pos or CD45RAposCD27pos), effector-like (Teff, CD45ROnegCCR7neg or CD45RAposCD27neg), central memory-like (Tcm, CD45ROposCCR7pos or CD45RAnegCD27pos), effector memory-like (Tem, CD45ROposCCR7neg or CD45RAnegCD27neg). For Anti-EGFR 4-1BB-CAR or sgRNA expression, respective reporter genes EGFP and mCherry fluorescence signals were analyzed. Mitochondrial mass was assessed by incubating cells with 10 nM MitoTracker Deep Red (Thermo Fisher M22426) for 20/30 min at 37 °C, followed by washing and analysis. All analyses were performed with a BD Fortessa (FACSDiva software v6.1.3).

### Correlation study on bioluminescence and infiltrating CAR-T cells % data

At the experiment endpoint, tumor charge assessed by bioluminescence and CAR-T cell abundance determined by flow cytometry were analyzed for correlation. Tumor charge confidence intervals for each point were used for goodness-of-fit assessment, defined by the standard deviation of photon rates (oscillating during measurement). After testing simple models, the best fit, evaluated by log-likelihood maximization, was a power law:

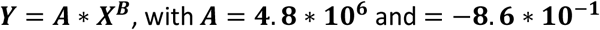

Optimization assumed Gaussian-like noise to avoid penalizing data-points with large error bars. This convention is widely used in counting experiments and justified in the literature [92]. No systematic error in measurements was considered.

### SgRNA readout by deep sequencing

The sgRNA library readout was performed using PCR amplicon sequencing. Genomic DNA was PCR amplified with adapters P5 and P7 bound custom primers (Eurofins) targeting a 350 bp vector region containing the sgRNA cassette, using adapters P5 and P7.

Adapter-Forward: ACACTCTTTCCCTACACGACGCTCTTCCGATCT-TTGCATATACGATACAAGGCTGT Adapter-Reverse: GACTGGAGTTCAGACGTGTGCTCTTCCGATCT-ACAAAGTTGGAATTCGGTACCTC PCR was carried out with Platinum™ SuperFi II Master Mix (ThermoFisher, 12368010) and 5% DMSO, with the following cycling conditions: 98°C for 8 minutes, 35 cycles of (98°C for 30 s, 58°C for 30 s, 72°C for 1 min), and 72°C for 10 minutes. Each reaction used 2 μg gDNA, with 3 independent reactions per sample. PCR products were pooled, gel-purified with a 2% agarose gel and FastPure Gel DNA Extraction Kit (Vazyme DC301-01), then normalized. The samples were sequenced by Eurofins Genomics using the NGSelect Amplicon Illumina MiSeq service, producing 60K reads per sample with basic quality control trimming of FASTQ files.

### MAGeCK analysis of gene KO subpopulations enrichment

For MAGeCK analysis, FASTQ files were processed with the MAGeCK-count algorithm to generate count tables, which were then input into the MAGeCK-test algorithm. (https://sourceforge.net/p/MAGeCK/wiki/Home/) The <mean= setting was used for LFC calculation, and normalization was set to <total= with non-targeting control sgRNAs for RRA null distribution, except for of subcutaneous tumor and lung samples from this experiment, where all gRNAs were used for normalization due to the extremality of data in these settings, allowing a more conservative but more effective calculation of p-values. LFC data were corrected to the mean LFC of non-targeting gRNAs, and ranking was done with MAGeCK RRA for most analyses. In the case of subcutaneous tumor and lung samples from this experiment, LFC-based ranking was applied, again to correct for the extremality of data. P-values and FDR values indicated for screening experiments are reported as calculated by the MAGeCK algorithm.

### Inducible cell death experiment

iCASP9-expressing CAR-T cells were stimulated with anti-CD3/CD28 TransAct (Miltenyi, 130-111-160) at 1:200 concentration for 48 hours. Cells were reseeded at 1×10v cells/ml and treated with 5 nM AP20187 (MedChemExpress HY-13992) or vehicle DMSO solution, in fresh medium. Cell survival was assessed at different time points by Fixable Live/Dead Blue (Thermofisher, L23105) staining, correcting for the actual number of cells when reductions were observed compared to time 0.

### CyTOF staining and data analysis

Frozen samples were thawed, washed in PBS with 0.5% BSA (BSA; Sigma-Aldrich, A7030), 0.02% sodium azide, and 15 µg/mL DNase, and stained with a surface antibody cocktail in PBS containing 0.5% BSA and 0.02% sodium azide for 15 minutes at RT (see Supplementary Table X for antibody details). Viability was assessed by incubating with 5 µM cisplatin for 5 minutes. After 2 washing steps, cells were permeabilized with Cyto-Fast Fix/Perm Solution (BioLegend) for 30 minutes at RT, stained intracellularly with anti-Perforin, anti-Granzyme A, anti-Granzyme B, anti-Granzyme K, and anti-CTLA-4 antibodies for 30 minutes at RT in permeabilization buffer, and fixed in 2% PFA overnight. DNA was stained as described previously. [93] Antibodies were obtained from suppliers listed in Supplementary Table 1 and conjugated following Standard Biotools’ protocol. CyTOF acquisition was conducted as previously described [94], and zero values were randomized using a custom R script, consistent with previous CyTOF software defaults. UMAP analysis was performed using custom R scripts with the “flowCore” and “uwot” packages as previously described, [95] and data transformation applied with the “logicleTransform” function from “flowCore” (parameters: w = 0.25, t = 16,409, m = 4.5, a = 0). For heatmap visualization, median intensity values were transformed using a logical data scale, following the approach described by Moore and Parks[96]. A four-color scale was used: black–blue (low), green–yellow (intermediate), and red (high) expression values

### *In vitro* restimulation assays

For in vitro screening restimulation assays, 5 × 10u mCherry+Luciferase+ A549 cells were seeded in 12-well plates (Corning, 351143). After 24 hours, plates were irradiated with 50 Gy using a BIOBEAM GM 2000 (Ekert and Ziegler), the medium was changed to Human T cell medium without interleukins, and lentiviral-modified CAR-T cells were added at a 2:1 effector-to-target ratio, 3 biological replicates were maintained for each donor in each condition. In the heavy restimulation assay, cocultures were refreshed every 5 days with medium changes every 3-5 days for 35 days. In the mild restimulation assay, cocultures were renewed every 10 days for 35 days. In both experiments cocultures were downsized to smaller wells when needed due to CAR-T cells population contraction, maintaining corresponding cells density. For the 14-day proliferation assay, CAR-T cells were cultured without tumor cells and restimulated every 5 days with anti-CD3/CD28 TransAct (Miltenyi 130-111-160) for 48 hours. After 14 days, cells were harvested for DNA extraction and compared to cells from the heavy restimulation assay at 14 days.

### Flow Cytometric Sorting of High and low MitoTracker Red-Expressing Cells

For sorting based on mitochondrial mass, cells were incubated with 10 nM MitoTracker Deep Red (Thermo Fisher M22426) at 37°C for 20-30 minutes, then washed and resuspended in sorting buffer (PBS with 2% FBS and 1 mM EDTA). Cell sorting was performed on a BD FACSAria™ III, adjusting FSC and SSC to exclude debris and doublets. Cells were gated by fluorescence intensity, and the highest (MitoTracker-High) and lowest (MitoTracker-Low) 10% populations were sorted into RPMI-1640 with 20% FBS. Sorted cells were pelleted, frozen, and used for DNA extraction to analyse gene-KO subpopulation frequency and mtDNA quantification.

### mtDNA quantification by qRT-PCR

Genomic DNA was extracted using the FastPure DNA Isolation Kit (Vazyme, DC112-01) and purified with silica columns. qRT-PCR was performed on a LightCycler® 480 system (Roche) with the SYBR Green I Master kit (Roche, 04707516001), with three replicates per sample. Analysis was performed with LightCycler® 480 Software (version 1.5.1.62) Primers used: ND1-Fw 5’-CCCTACTTCTAACCTCCCTGTTCTTAT-3’, ND1-Rv 5’-CATAGGAGGTGTATGAGTTGGTCGTA-3’, ND5-Fw 5’-ATTTTATTTCTCCAACATACTCGGATT-3’, ND5-Rv 5’-GGGCAGGTTTTGGCTCGTA-3’, 18S-Fw 5’-AGTCGGAGGTTCGAAGACGAT-3’, 18S-Rv 5’-GCGGGTCATGGGAATAACG-3’. Mitochondrial DNA levels were calculated using the ΔΔCt method with 18S rRNA as the reference gene.

### In vitro motility assay with collagen gel and Analysis of T cell motility

T cells were stained with 100 nM Calcein Green (Invitrogen C34852) or Calcein Red Orange (Invitrogen C34851) for 20 min at 37 °C in DPBS, washed, and resuspended in 50 µl motility medium at 20 × 10v cells/ml. T cells were mixed with 100 ul of 2 mg/ml collagen solution (PureCol Bovine Collagen, CellSystem 5005), loaded into ibiTreat μ-Slide VI 0.4 (Ibidi 80606), and incubated at 37 °C for 1 h for collagen polymerization. T cell motility was recorded using a Nikon epifluorescence microscope, equipped with 37 °C heated-chamber, with ORCA-Flash 4.0 camera (C11440 Hamamatsu) and Metamorph software (version 7.10.2.240). Images (15 per condition, 30 s interval, 4/5 z-planes, z-distance of 15 µm) at 3 different stages were acquired using a 20x objective (binning rate 2). In most cases, T cells from two donors were stained with Calcein Green and Red, mixed, and added to collagen.

Motility analysis was done using TrackMate78 (ImageJ plugin, v.7.11.1) in 3D, LoG detector was used to identify spots (quality and maximal intensity filters applied) and Simple LAP Tracker to identify tracks (linking max distance 20 µm, closing-gap max distance 20 µm, frame gap 3) Only tracks with 10+ spots were analyzed. Motility Score was calculated as (% cells moving >10 µm) * (mean speed [µm/min] of moving cells). Analysis used original 3D images, considering z-axis displacement.

### High-resolution respirometry with Oroboros device

The oxygen consumption rate (OCR) was measured using a 2 mL Oxygraph-2k respirometer (Oroboros). The electrode was calibrated at 37 °C, 100% and 0% oxygen, then CD8+ T cells (2 mL at 2.5 × 10v cells/mL) were added to the chamber. O2 consumption flux was measured in pmol/s. Oligomycin (0.5 μg/mL) was added to block complex V, followed by increasing concentrations of carbonyl cyanide m-chlorophenyl hydrazone (CCCP) to assess maximum respiratory capacity. Potassium cyanide (KCN) was added to measure non-mitochondrial respiration.

### Statistical analysis

Data are shown as mean ± SEM or box-and-whisker plots. Statistical comparisons for normally distributed groups were performed using paired or unpaired two-tailed Student’s t-test (for two-group comparisons) or One-way/Two-way ANOVA. For non-normally distributed data, the Wilcoxon Signed Rank Test (paired) or ANOVA on Ranks was used. Post hoc tests were applied for pairwise comparisons. Analyses were conducted using GraphPad (v8).

### Graphics

All schematic representations were designed with Biorender.com

## ACKNOWLEDGEMENTS

This work was supported by INSERM, the French Ligue Nationale Contre le Cancer (LNCC), the Cochin Institute (POC 2020 and 2025, PIC 2025), Inserm Transfert (CoPOC), Universite Paris Cite (Ami-Stratex 2024), and the Laboratory of Excellence GR-EX. M.F. and D.A. were supported by PhD fellowships from the Ligue Nationale Contre le Cancer and the China Scholarship Council (CSC), respectively. M.F. also received support from the <EUR G.E.N.E Fellowship= during his Master’s internship.

We are extremely grateful to the members of the following platforms and facilities: the Cochin Institute Imaging Facility (IMAG’IC); the Cytometry and Immunobiology Platform (CYBIO), with special thanks to Youcef Hadjou and Sougnaya Many; the Histology, Immunostaining, and Microdissection Facility (HistIM), especially Maryline Favier, Fabiola Ely-Marius, and Rachel Onifarasoaniaina; and the In Vivo Imaging Facility (PIV), with special thanks to Gilles Renault, Isabelle Lagoutte, Franck Lager, Pierre Monbernard, and Giovanna Bibaki.

We thank Pierre de la Grange (Genosplice) and Olivier Tenaillon for helpful discussions on the interpretation of statistical analyses, and Jaques Dutrieux for his help on the set up of the PCR protocols. We are also grateful to Concetta Quintarelli, Simona Manni, Beatrice Mezzopra, Julien Sage, Isabelle Munoz, and Marianne Mangeney for their valuable scientific input.

Finally, we thank all members of the <Cancer Immunotherapy and Cell Reprogramming= and <Cancer and Immune Response= teams, especially Alice Machado, Marwan Fontaine, Jeane Pitiot and Irena Rajnpreht, for their daily support and constructive scientific feedback.

## AUTHOR CONTRIBUTIONS

Conceptualization & Writing – Original Draft: M.F. and F.P. Funding Acquisition and Resources: F.P., N.B., and E.D.

Investigation: M.F., D.A., C.C., L.S., L.A., Y.S., M.C.-A.G., A.V., A.V., M.M., J.M., N.T., N.B., and E.D.

Writing – Review & Editing: N.T., J.M., N.B., and E.D. All authors read and approved the final manuscript.

## MATERIALS AND CORRESPONDENCE

Frederic Pendino and Mattia Fumagalli

---

**Supplementary Figure 1.**
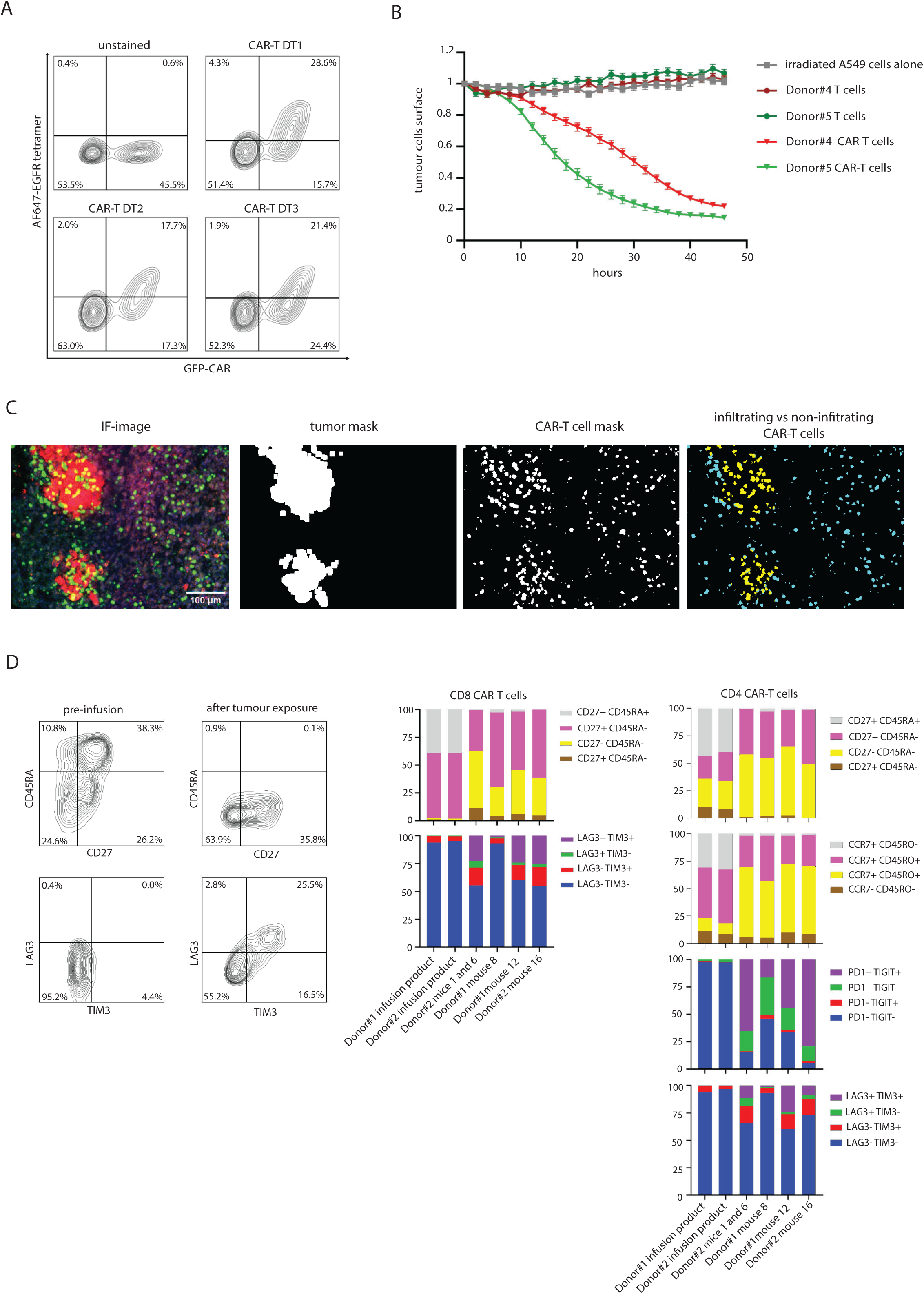
**Validation of CAR-T cells specificity and of the orthotopic lung tumour model**: **(A)** CAR-T cells binding with biotin-EGFR recombinant protein conjugated in tetramers with A647-streptdadivin. **(B)** CAR-T cells mediated killing of A549 cells as assessed as reduction of mCherry+ area, measured by Incucyte imaging. **(D)** CAR-T cells infiltration in tumour islets quantification strategy, from left to right images of different masks used, and merged image of CAR-T cells colocalizing (yellow) or not (cyan) with the EpCAM mask. **(E)** Phenotypical characterisation of CAR-T cells by flow cytometry. We compared CAR-T cells from infusion products and CAR-T cells recovered from mice which received the 5*10˄6 dose after long term exposure to tumour *in vivo*.

**Supplementary figure 2.**
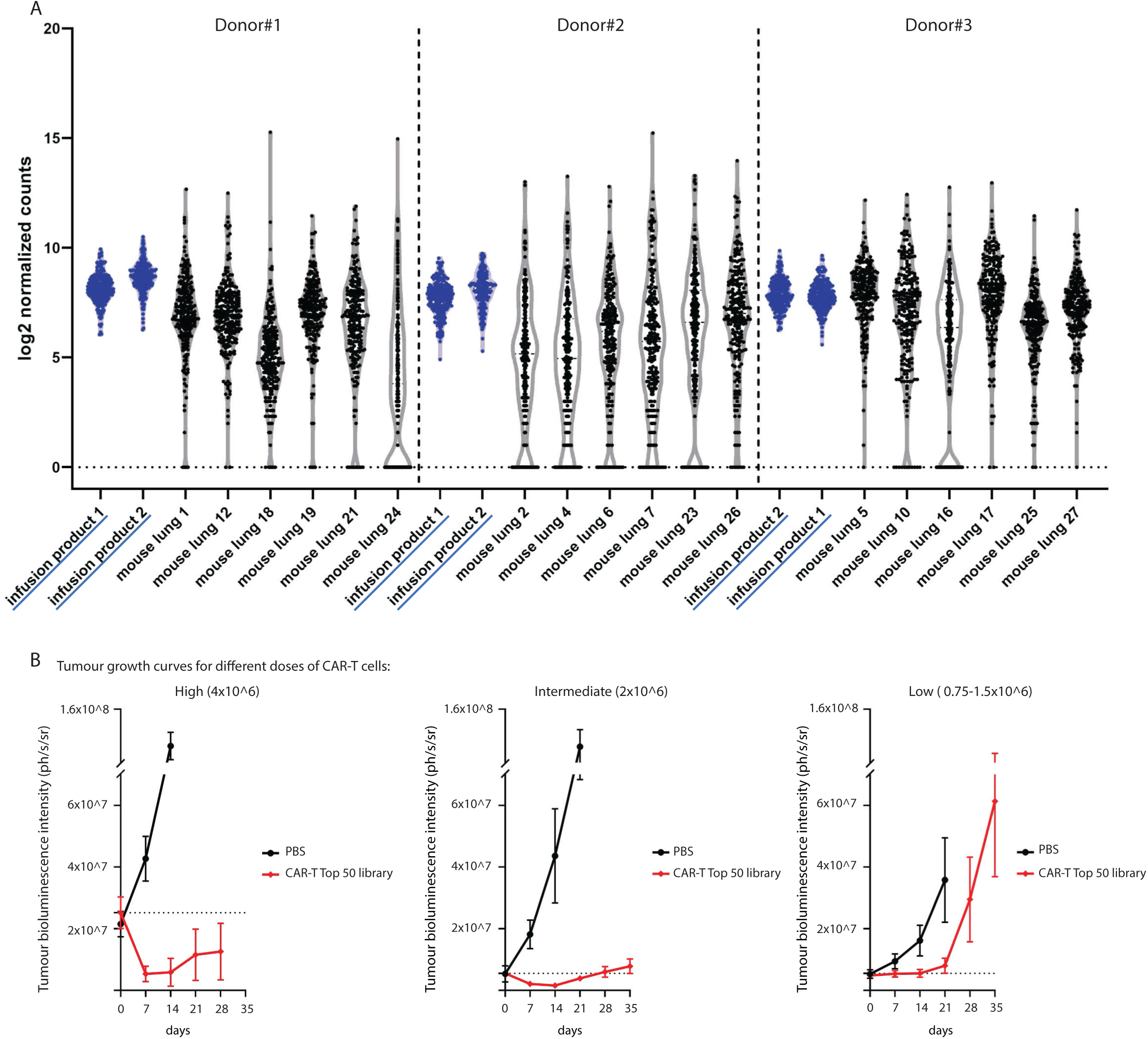
**Additional details on the *in vivo* screening assays**: (**A**) log2 counts of sgRNA in CAR-T cells from the three different donors, compared in mice lungs at the endpoint of the experiment vs in infusion products. For CAR-T cells from donor#1 data from mice 19 and 24, that died before the expected endpoint, (among 21 and 28 days) were included too. For CAR-T cells from donor#3, DNA sample from mouse 11 could not be treated. (**B**) Tumour growth curves as determined by bioluminescence for mice in different experiments on the same donor CAR-T cells, Donor#3. Mice first received 10*10˄6 A549 cells in the first experiment and 4*10˄6 In both the other ones. In the first experiment mice are treated either with excipient (n=5) or 4*10˄6 CAR-T modified with our Top 50 library lentivirus bank (n=5), in the second experiment they are treated either with excipient (n=5) or 2*10˄6 CAR-T modified with our Top 50 library lentivirus bank (n=5), in the third experiment they are treated either with excipient (n=5) or 0.75-1.5*10˄6 CAR-T modified with our Top 50 library lentivirus bank (respectively n=4 and n=4),

**Supplementary figure 3.**
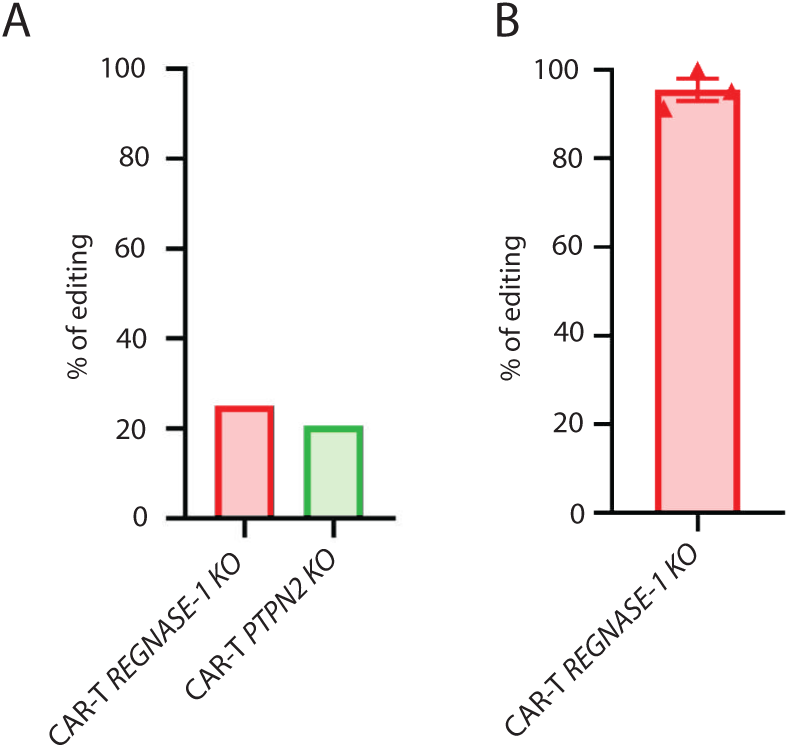
**Editing validation for *PTPN2* KO and *REGNASE-1* KO in different experiments**: (**A**) quantification of *REGNASE-1* and *PTPN2* gene editing in CAR-T cells used for the validation in vivo experiments, as evaluated by proportion of modified genomic sequences upon knock out, determined through sanger sequencing and deconvolution analysis of chromatograms files. (**B**) Quantification of *REGNASE-1* gene editing in CAR-T cells used for the in vitro suicide gene experiment, as evaluated by proportion of modified genomic sequences upon knock out, determined through sanger sequencing and deconvolution analysis of chromatograms files. (n=3) To maximize editing efficiency, we used a double gRNA approach with recently described highly effective *REGNASE-1* targeting sgRNA sequences.

**Supplementary Figure 4.**
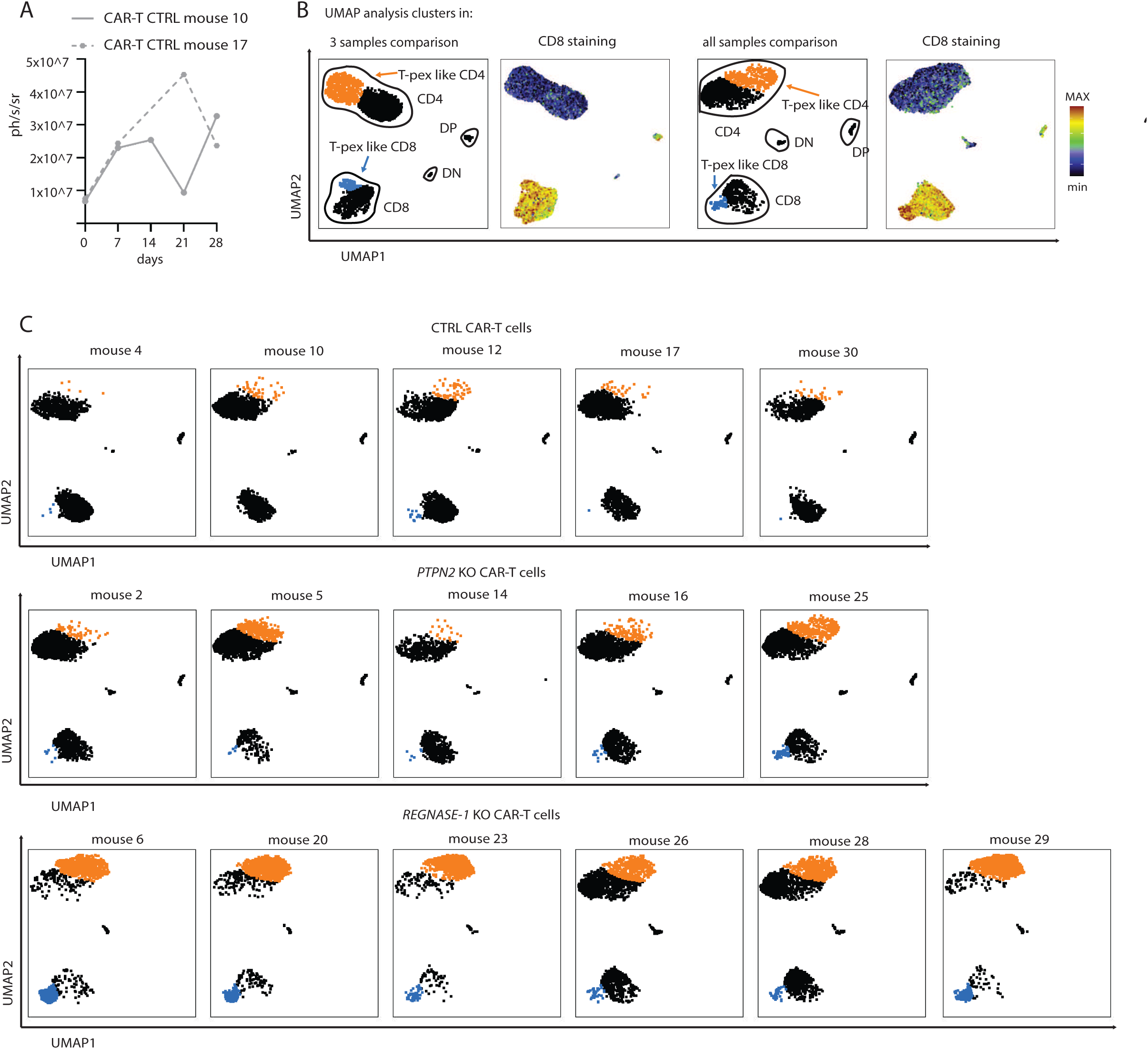
Detailed sample-by-sample clustering analysis of CyTOF data: (**A**) Tumour growth curve in mice 10 and 17, these mice were excluded from the tumour control comparison experiment, because of excessive tumour burden. These mice were treated with a higher dose of CTRL CAR-T cells, 2*10˄6 and were used as additional controls exclusively for the CyTOF experiment. (**B**) Comparison of UMAP analysis of CyTOF data from CAR-T cells in lung tumour samples at the endpoint of the experiment, either confronting clustering in the analysis with three exemplary samples, as in the main figure 5, or including all samples in the analysis. (**C**) UMAP analysis of CyTOF data from CAR-T cells in lung tumour samples at the endpoint of the experiment, in this analysis all samples were included.

**Supplementary figure 5.**
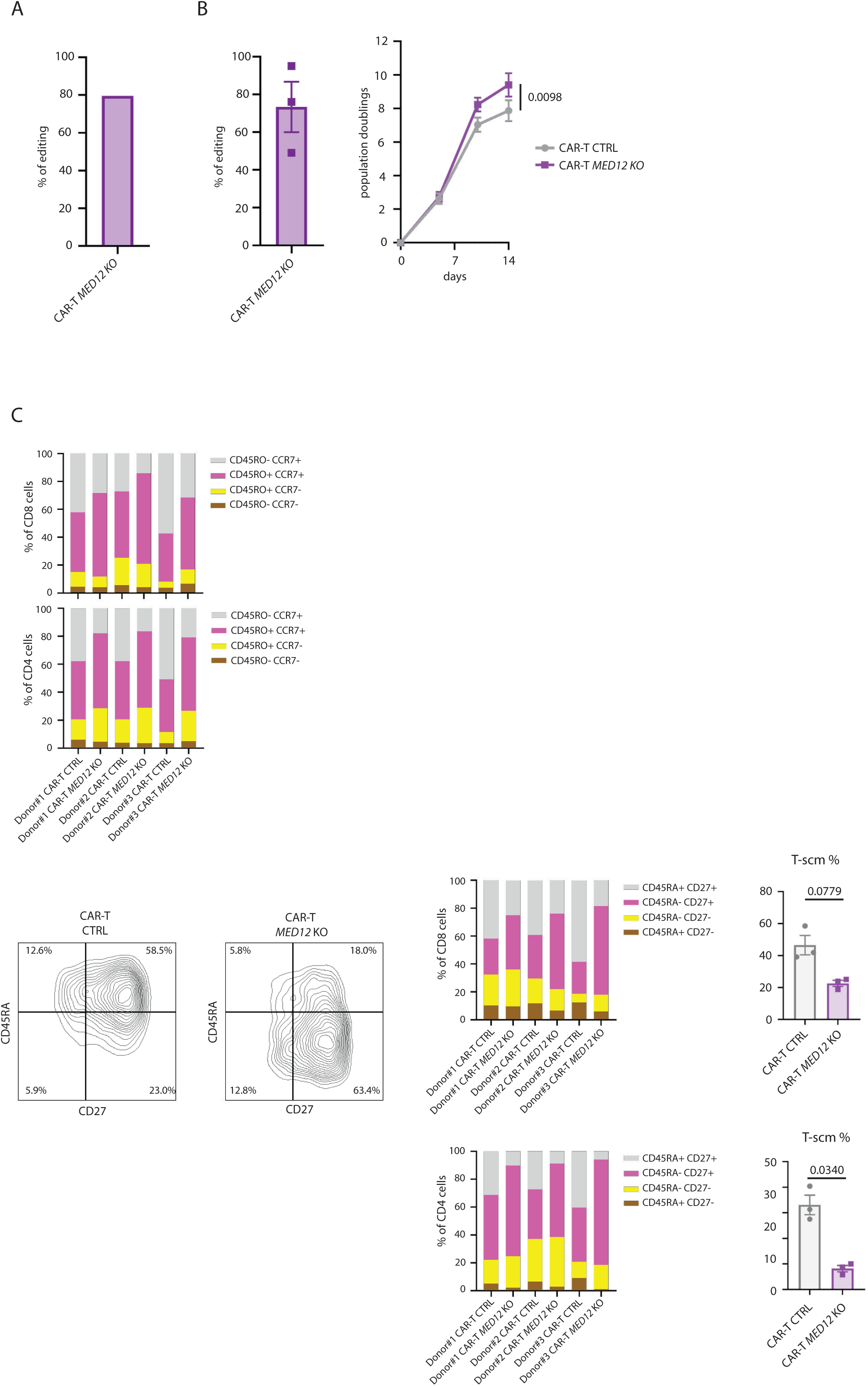
**Validation of *MED12* editing in different experiments and additional characterisation of *MED12* KO CAR-T cells**: **(A)** quantification of *MED12* gene editing in CAR-T cells used for the validation in vivo experiment, as evaluated by proportion of modified genomic sequences upon knock out, determined through sanger sequencing and deconvolution analysis of chromatograms files. **(B)** (left) Quantification of *MED12* gene editing in CAR-T cells used for in vitro characterisation experiments, as evaluated by proportion of modified genomic sequences upon knock out, determined through sanger sequencing and deconvolution analysis of chromatograms files. (right) *In vitro* proliferation during production phase for *MED12* KO CAR-T cells as compared to CTRL CAR-T cells (n=3) **(D)** Flow cytometry-based differentiation profiling for *MED12* KO CAR-T cells as compared to CTRL CAR-T cells at the endpoint of production phase. (n=3 per group) T-scm (T stem cell memory) are here defined as CD45RA+ and CD27+ t-cells, to confirm data from the main figure by a different staining. P-values are calculated with unpaired t-tests.

**Supplementary figure 6.**
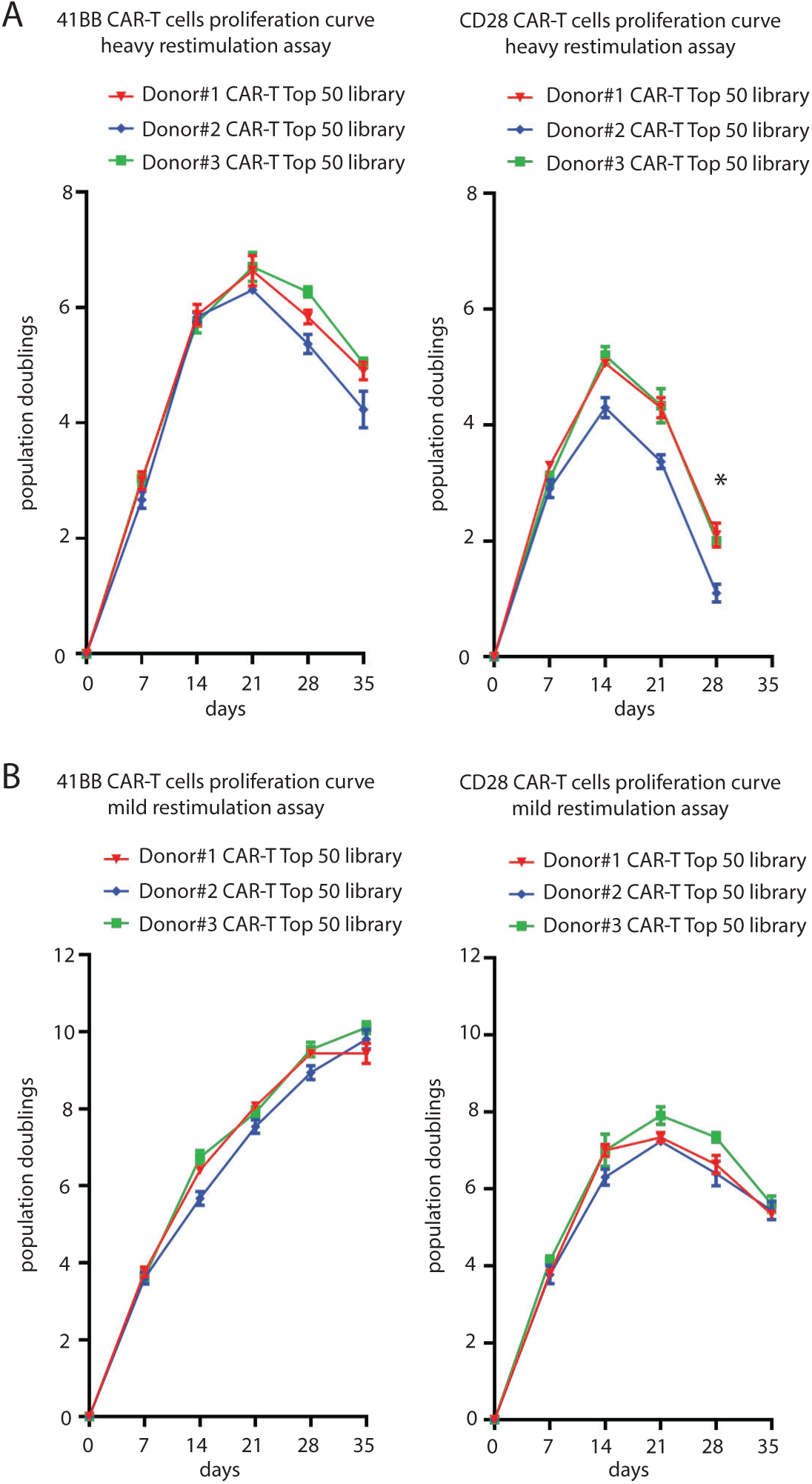
**41BB-CAR-T and CD28-CAR-T cells proliferation curves in restimulation assays**: **(A)** 41BB-CAR-T and CD28-CAR-T cells proliferation curve, expressed in population doublings, in the heavy restimulation assay. Numbers are determined by Trypan Blue corrected cell counting, we cultured 3 replicates for each of 3 donors derived CAR-T cells Donor#1, Donor#2, Donor#3. For the CD28 condition, (* at 28 days we could not detect enough live cells to relaunch the co-culture.) **(B)** 41BB-CAR-T and CD28-CAR-T cells proliferation curve, expressed in population doublings, in the mild restimulation assay. Numbers are determined by Trypan Blue corrected cell counting, we cultured 3 replicates for each of 3 donors derived CAR-T cells Donor#1, Donor#2, Donor#3.

**Supplementary Fgure 7.**
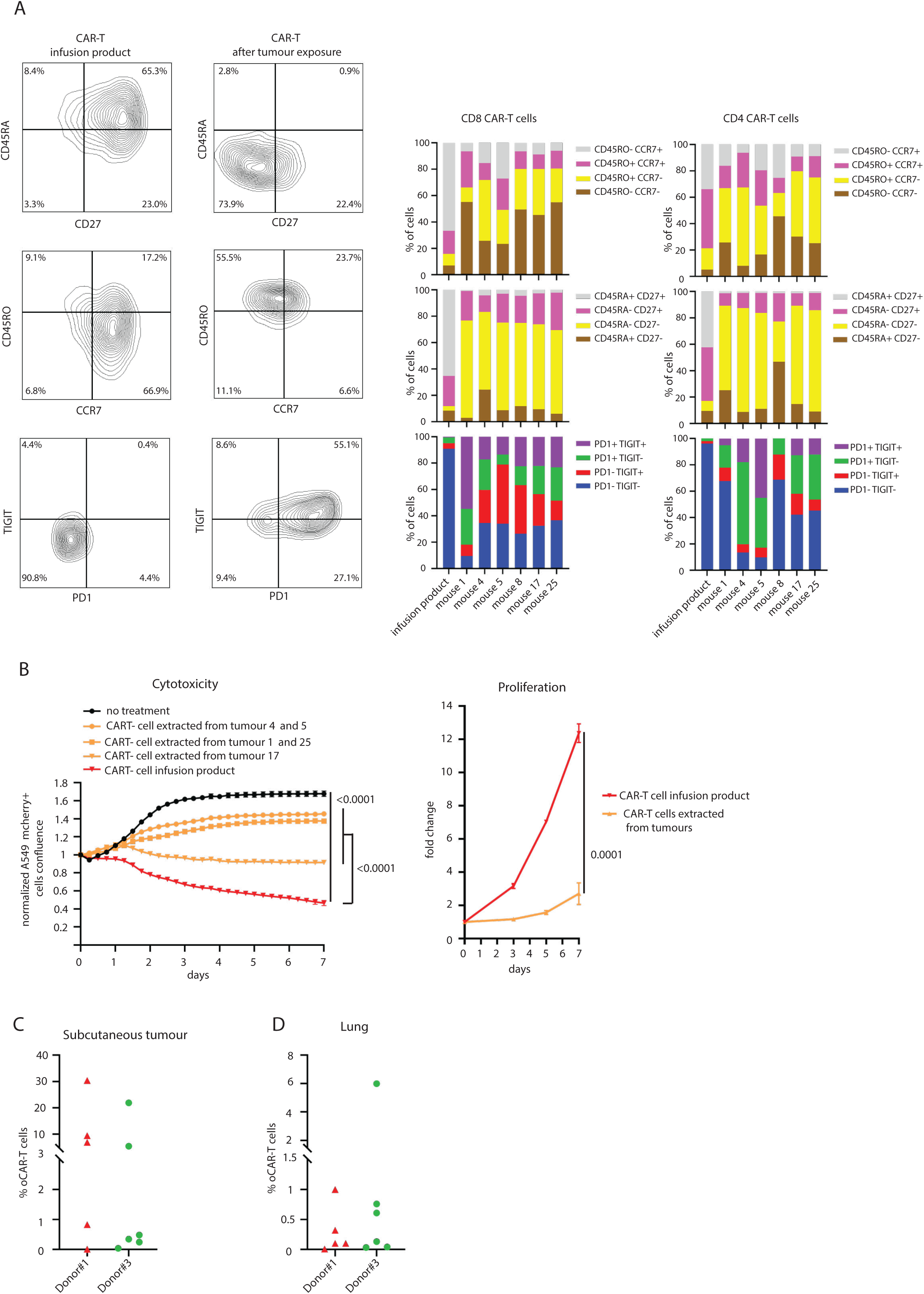
**Characterization of the A549 subcutaneous tumour model**: **(A)** Phenotypical characterisation of CAR-T cells by now cytometry. We compared CAR-T cells from infusion products and CAR-T cells recovered from mice subcutaneous tumours which received 5*10˄6 CAR-T cells, after long term exposure to tumour *in vivo*. **(B)** Functional characterisation of compared CAR-T cells from infusion products and CAR-T cells recovered from mice. Cytotoxic capacity was assessed by calculating surface reduction of mCherry+ tumour cells when co-cultured with CAR-T cells. Proliferation capacity was assessed by cell counting upon 7 days after anti-CD3/C28 stimulation. P-values are calculated by two-ways ANOVA test, results for onal time point comparisons are shown. **(C)** CAR-T cells detection in mice subcutaneous tumours and **(D)** lungs by now cytometry, for Donor#1 (n=5) and Donor#3 (n=6)

**Supplementary Fgure 8.**
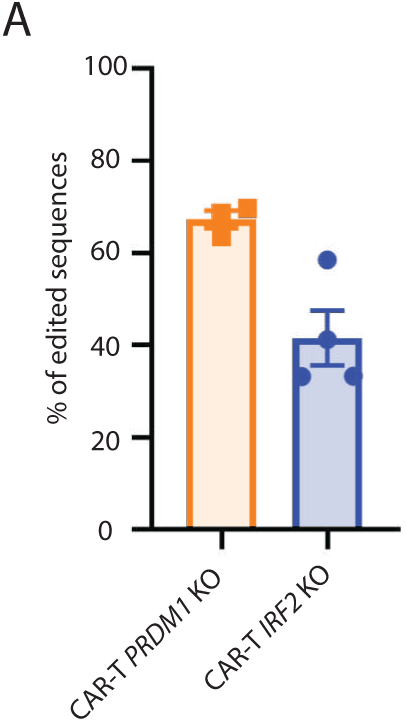
***PRDM1* and *IRF2* gene editing validation**: **(A)** Quantification of *PRDM1* and *IRF2* gene editing in CAR-T cells used for all the in vitro experiments, as evaluated by proportion of modified genomic sequences upon knock out, determined through sanger sequencing and deconvolution analysis of chromatograms Files. (n=4 for group)

